# Bayesian metamodeling of complex biological systems across varying representations

**DOI:** 10.1101/2021.03.29.437574

**Authors:** Barak Raveh, Liping Sun, Kate L. White, Tanmoy Sanyal, Jeremy Tempkin, Dongqing Zheng, Kala Bharat, Jitin Singla, ChenXi Wang, Jihui Zhao, Angdi Li, Nicholas A. Graham, Carl Kesselman, Raymond C. Stevens, Andrej Sali

## Abstract

Comprehensive modeling of a whole cell requires an integration of vast amounts of information on various aspects of the cell and its parts. To divide-and-conquer this task, we introduce Bayesian metamodeling, a general approach to modeling complex systems by integrating a collection of heterogeneous input models. Each input model can in principle be based on any type of data and can describe a different aspect of the modeled system using any mathematical representation, scale, and level of granularity. These input models are (i) converted to a standardized statistical representation relying on Probabilistic Graphical Models, (ii) coupled by modeling their mutual relations with the physical world, and (iii) finally harmonized with respect to each other. To illustrate Bayesian metamodeling, we provide a proof-of-principle metamodel of glucose-stimulated insulin secretion by human pancreatic ß-cells. The input models include a coarse-grained spatiotemporal simulation of insulin vesicle trafficking, docking, and exocytosis; a molecular network model of glucose-stimulated insulin secretion signaling; a network model of insulin metabolism; a structural model of glucagon-like peptide-1 receptor activation; a linear model of a pancreatic cell population; and ordinary differential equations for systemic postprandial insulin response. Metamodeling benefits from decentralized computing, while often producing a more accurate, precise, and complete model that contextualizes input models as well as resolves conflicting information. We anticipate Bayesian metamodeling will facilitate collaborative science by providing a framework for sharing expertise, resources, data, and models, as exemplified by the Pancreatic ß-Cell Consortium.

**Significance Statement:** Cells are the basic units of life, yet their architecture and function remain to be fully characterized. This work describes Bayesian metamodeling, a modeling approach that divides-and-conquers a large problem of modeling numerous aspects of the cell into computing a number of smaller models of different types, followed by assembling these models into a complete map of the cell. Metamodeling enables a facile collaboration of multiple research groups and communities, thus maximizing the sharing of expertise, resources, data, and models. A proof-of-principle is provided by a model of glucose-stimulated insulin secretion produced by the Pancreatic ß-Cell Consortium.

## Introduction

### Modeling of the cell

Cells are the basic structural and functional units of life (1). Different aspects of the cell have been studied extensively, including experimentally, computationally, and theoretically. As is the case for any model, a cell model is expected to provide more information about the cell than any of the input information used for its construction. In particular, the model should rationalize known facts and make testable predictions. We consider a desired cell model and its construction by discussing a progression from an impractical atomic model, an impractical integrative model, actual current models, and finally culminating in the modeling approach proposed here.

### Impractical physical modeling of the cell

Hypothetically, a most precise and complete model of the physical cell structure specifies trajectories for each of its atoms over its lifespan. Such a model could in principle be obtained from molecular dynamics simulations (2, 3). In practice, however, computing accurate trajectories for ∼10^14^ atoms over days or longer is limited by inaccurate molecular mechanics force fields, slow computers with insufficient memory, as well as lack of sufficient knowledge about the starting state and environment. Moreover, even if such a model could be computed, it still would not abstract all cellular properties of interest, such as molecular signalling networks.

### Recalcitrant integrative modeling of the cell

To attempt to address these challenges, we could adopt an integrative approach. Integrative modeling is defined as modeling that uses multiple types of information about the modeled system, be it from different experiments or prior models (4). It is motivated by the resulting increase in accuracy, precision, and completeness of a model. Integrative modeling is particularly effective for modeling complex biological systems, for which no single experimental or theoretical approach can provide all needed information. For example, structures of large macromolecular assemblies recalcitrant to traditional structural biology methods have been determined by integrative structure determination (5). Integrative modeling of the cell could rely on a multiscale representation and multimodal experimental data, in addition to the first principle of physics (4). In practice, however, even integrative modeling of all aspects of the entire cell is not feasible at this time, due to insufficient data and computing power as well as limitations of existing integrative modeling methods.

### Current models of aspects of the cell

Although an accurate, precise, and complete model of the cell cannot yet be computed, it is possible to model some aspects of the cell or its parts with useful accuracy and precision. Most of these models rely on a single type of representation of the cell; for example, spatiotemporal (6), ordinary differential equation (ODE) (7), and flux balance analysis representations (8). In addition to whole-cell models, there are a myriad of models of different parts of the cell, too numerous to review here. These models may provide a useful starting point for whole-cell modeling, due to their encoding of expertise, data, and computing used to produce them. However, no general approach yet exists for combining different kinds of models, although steps in this direction have been made (Discussion) (9–13).

### Bayesian metamodeling of the cell

Here, we propose a divide-and-conquer modeling approach that integrates input models of varied representations into a metamodel. Metamodeling can be seen as a special case of integrative modeling in which the focus is on integrating prior models instead of data (4). The large problem of computing an integrative model of the cell is broken into a number of smaller modeling problems corresponding to computing models of some aspects of some parts of the cell. Each such input model may be informed by different subsets of available data, relying on its distinct model representation at any scale and level of granularity. Metamodeling then proceeds by assembling and harmonizing the input models into a complete map of the cell. Here, the input models are harmonized through a Bayesian statistical model of their relations with each other and/or the physical world. This Bayesian approach enables us to update our “beliefs” in the distribution of model variables (including best single-value estimates and their uncertainties), given information provided by all input models. By shifting the focus from data integration to model integration, Bayesian metamodeling facilitates the sharing of data, computational resources, expertise in diverse fields, and already existing models of the cell and its parts.

### Proof-of-principle: Prototype metamodel of glucose-stimulated insulin secretion (GSIS)

The Pancreatic ß-Cell Consortium (pbcconsortium.org) brought together research groups in biology, chemistry, physics, mathematics, computer science, and the digital arts (14). The consortium provides a nurturing environment for developing methods for whole-cell modeling. For developing the method of metamodeling, we narrowed our focus on glucose-stimulated insulin secretion (GSIS) (15), one of the key functions of the ß-cell (Fig. 1). Insulin secretion encompasses many of the complexities of the whole cell, including aspects that are best described using different types of models at different scales, thus providing a useful testing ground for Bayesian metamodeling of the cell.

**Figure 1.**
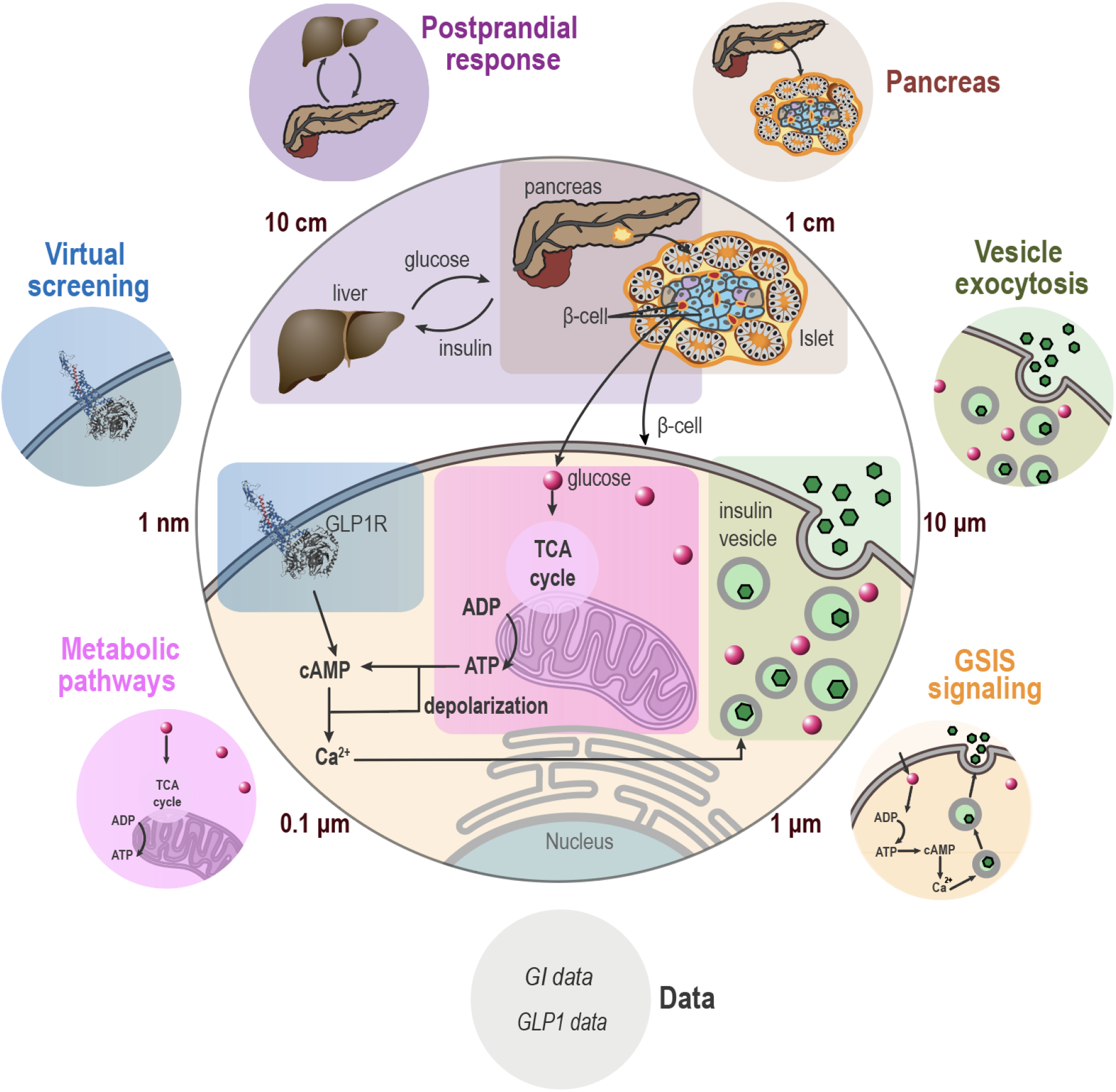
Metamodeling of glucose-stimulated insulin secretion (GSIS). Eight input models, including two data models, describe different aspects of GSIS (Table 1). They are represented by small circles with different background colors. These input models are integrated into a single metamodel of GSIS, indicated by a large grey circle in the center.

## Results

### Definitions

We are using a number of common terms that may have different definitions in different fields. Thus, we begin by defining our usage here. We are working in the Bayesian framework that estimates a model based on data and prior information (16). Thus, a *model* is the joint posterior probability density function (PDF) over the model variables. We distinguish among three kinds of model variables. First, *free parameters* (*i*.*e*., degrees of freedom) are quantities that are fit to input information (*e*.*g*., the coefficients of a polynomial model and atomic coordinates of a protein structure model). Second, *independent variables* (*i*.*e*., regressors, features, or inputs) are quantities whose values are supplied when evaluating a model (*e*.*g*., the abscissa of a polynomial model). Third, *dependent variables* (*i*.*e*., response variables, regressands, outcomes, labels, predictions, or outputs) are quantities whose values are computed when evaluating a model (*e*.*g*., the ordinate of a polynomial model). As an aside, *fixed parameters* (*i*.*e*., constants or hyperparameters) are quantities whose values are defined and fixed (*e*.*g*., stereochemistry parameters in protein structure modeling). *Systematic error* of a model (variable) is the mean difference between the model (variable) and the “ground truth”. *Random error* of a model (variable) is the spread (*e*.*g*., standard deviation, standard error of the mean, and entropy) of the model (variable). While the ground truth is never known, systematic error can still be approximated by the difference between the model and an independent reference that represents the ground truth as closely as possible (a gold standard). Accuracy and precision (uncertainty) are equivalent to systematic error and random error, respectively, except they increase as their counterparts decrease. Ideally, accuracy is approximated by precision. Metamodeling couples and harmonizes all input models by updating the PDFs of their free parameters.

### The input models

Information for GSIS metamodeling is provided by eight input models (Fig. 1; Table 1; SI Appendix: Supplementary Text 1). The models have been selected to cover a diverse range of representations, spatiotemporal scales, and data. They include a coarse-grained spatiotemporal simulation of insulin vesicle exocytosis, a molecular network model of GSIS signaling, a network model of insulin metabolism, an atomic structural model of glucagon-like peptide-1 receptor (GLP1R) activation, a linear model of the pancreatic cell population, ODEs for systemic postprandial insulin response, synthetic data on glucose intake (glucose intake data model), and synthetic data on GLP1 and GLP1 analogue levels (GLP1 data model).

**Table 1:** The input models for GSIS metamodeling.

| Model (abbreviation) | Representation | Description | Scale (spatial, temporal) | Granularity (spatial, temporal) | Experimental data and prior information | Ref. |
| --- | --- | --- | --- | --- | --- | --- |
| Postprandial response (PR) | Ordinary differential equations | Model of systemic postprandial change in plasma insulin and glucose levels after a meal | Body<br>$10^3$ sec | Organ<br>$10^1$ sec<sup>‡</sup> | Insulin, glucose measurements | (20)<br>1.1* |
| Pancreas (Pa) | Linear equations | Model of insulin secretion by the entire population of β-cells in the pancreas | Organ<br>N/A | Cell<br>N/A | Microscopic examination and morphometric measurements for quantifying islets in a pancreas (85); electrical tomography and morphometric analysis for quantifying β-cells in an islet (86). | 1.2* |
| Vesicle exocytosis (VE) | Spatiotemporal | Coarse-grained Brownian dynamics simulation of insulin granule trafficking, docking, and exocytosis | Cell<br>$10^{-1}$ sec | Molecule/Granule<br>$10^{-8}$ sec | Soft X-ray tomography (19) | (87)<br>1.3* |
| GSIS signaling (Sg) | Network / Linear ordinary differential equations | Model of signaling pathway of glucose-stimulated insulin secretion | Cell<br>$10^2$ sec | Molecule<br>N/A | KEGG pathways (88); fluorescence imaging and Förster-resonance energy transfer microscopy (FRET) (89–92) | 1.4* |
| Insulin metabolism (Mb) | Network | Model of cellular metabolic pathways upregulation or downregulation under different treatment conditions | Cell<br>$10^2$ sec | Molecular concentration<br>N/A | Proteomic/metabolomic screens, KEGG pathways (88) | 1.5* |
| Virtual screening of GLP1R (VS) | Spatial | List of GLP1R ligands ranked by their estimated activation of GLP1R, based on structure-based virtual screening of a library of GLP1 analogs | Macro-molecule<br>N/A | Atom<br>N/A | Virtual screening assay (93) based on structure-based modeling and X-ray crystallography | 1.6* |
| Glucose intake data (GI) | Time series | Rate of glucose intake after a meal | Body<br>$10^3$ sec | Organ<br>$10^1$ sec | Synthetic data derived from the glucose rate of appearance from the postprandial response model | 1.7* |
| GLP1 data (GL) | Synthetic data | GLP1R activation at different agonist levels | Macro-molecule<br>< sec | Atom<br>N/A | Synthetic data | 1.8* |

<sup>‡</sup> The model is continuous, but trained over data obtained at 1-minute resolution, with the precision on the order of seconds.
\* The model was computed here, based on prior publications, as described in SI Appendix: Supplementary Text 1.

### Bayesian metamodeling workflow

Given the input models, Bayesian metamodeling proceeds through three steps (Fig. 2): (i) conversion of the input models into surrogate probabilistic models; (ii) coupling of these surrogate models through subsets of statistically related variables; and (iii) backpropagation to update the original input models by computing the PDFs of free parameters for each input model in the context of all other input models. Thus, the output from metamodeling includes the joint PDF over all surrogate and reference variables (Step 2) as well as the updated input models (Step 3). We now describe each step in turn, both in general terms and by one or more examples.

**Figure 2.**
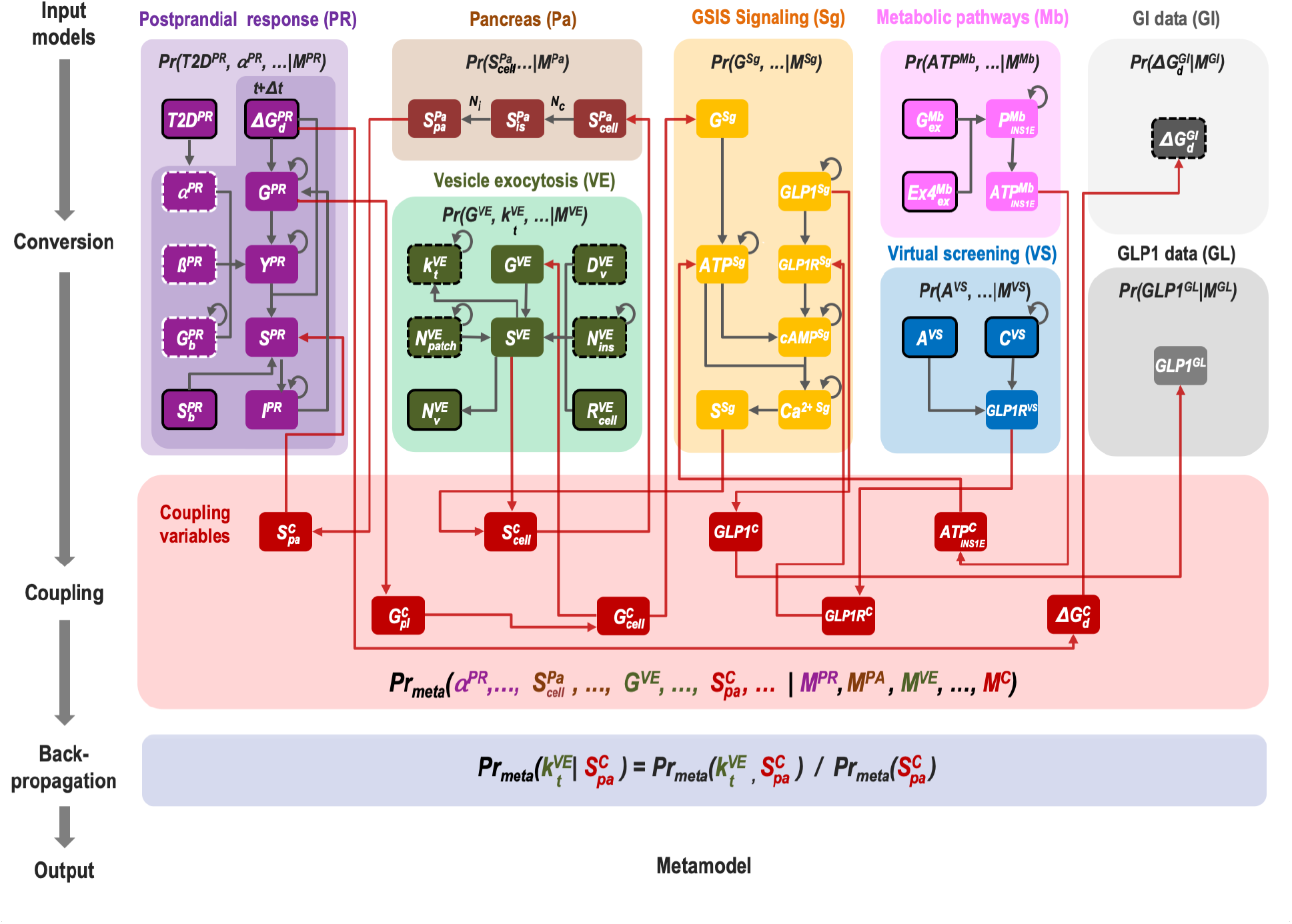
From input models to coupled surrogate models in a metamodel of GSIS. Nodes indicate variables and directed edges indicate probabilistic relations between a parent and child variable in a Bayesian network; a child variable is conditionally independent of any of its non-descendants, given the values of its parent variables (17). Each model and its variables are indicated by a specific color. Reference variables are in red, data variables are in grey, fixed parameters in the input models are encircled in white dashed lines, free parameters are encircled in black dashed lines, independent variables are encircled in continuous line, and dependent variables are not encircled. Grey edges are defined by the input models, whereas red edges are defined by the couplers. Self-loops indicate dependency on the value of the same variable in a previous time slice. Annotated variables and edges indicate examples discussed in the text.

### Step 1: Conversion of input models into surrogate probabilistic models

We first create a common representation for each input model by converting it into a corresponding probabilistic model. The probabilistic model is a surrogate for the original input model: it provides a probabilistic description of the input model, potentially simplifying it. Formally, the surrogate model specifies a PDF over some input model variables and any additional variables deemed necessary. This PDF encodes model uncertainty and statistical dependencies among its variables. Model uncertainty arises from insufficient information, imperfect modeling, and/or stochasticity of the system. Statistical dependencies are exemplified by the dependency between the values of independent and dependent variables, the effect of free parameter values on such dependency, and spurious correlations due to confounding factors or coincidence.

In principle, a surrogate model could be obtained by any approach for modeling statistical distributions, such as probabilistic graphical models (PGMs) (17) and various deep generative models (18). For the current proof-of-concept, we used Bayesian networks (BNs), which are a well-studied class of PGMs (17). BNs are often used for representing PDFs over many variables using a directed graph (network), with nodes and edges representing variables and conditional statistical dependencies, respectively. They are supported by efficient methods for Bayesian inference, parameter fitting, and learning of network topology from data. Finally, BNs include Dynamic Bayesian Networks (DBNs) that generalize both Hidden Markov Models and Kalman Filters, allowing us to model dynamic processes (17), such as vesicle exocytosis. In a nutshell, for a given input model, we tabulate dependent variables as a function of free parameters and independent variables, followed by manually constructing and parametrizing a surrogate PGM that approximates this table.

### Examples

We constructed a surrogate model for the vesicle exocytosis model (Figs. 1 and 2, green). Insulin vesicle exocytosis is described by spatiotemporal trajectories of insulin granules undergoing trafficking, docking, and exocytosis within a pancreatic β-cell over 200 milliseconds, following glucose stimulation (SI Appendix: Supplementary Text 1.3; SI Appendix: Movie S1) (19). A simplified cell representation incorporates the cell membrane, nucleus, hundreds of insulin vesicles, and thousands of glucose molecules. The trajectories of these components are computed using Brownian dynamics simulations restrained by various experimental data, including soft X-ray tomograms of the cell. The free parameters include parameters of the data-driven potential function and diffusion constants. The independent variables are the coordinates of the starting configuration. The dependent variables are millions of cell frames in a trajectory, each specifying coordinates of thousands of components. For practical reasons, the proof-of-principle surrogate model abstracts the billions of variables describing a trajectory by sampling it at a fraction of frames and including only a subset of variables for each sampled frame. Uncertainty in the values of the surrogate model variables reflects uncertainty in the corresponding input model.

As a second example, we constructed a surrogate model for the postprandial response model (20) (Figs. 1 and 2, purple; SI Appendix: Supplementary Text 1.1). Insulin and glucose fluxes through different body systems in the hours following a meal are described by ODEs. The variables of the postprandial response surrogate model include free parameters of the model ODEs (*i*.*e*., coefficients in ODEs) in either healthy or type 2 diabetic subjects; independent variables corresponding to the change in plasma glucose levels due to digestion; and dependent variables indicating predicted glucose and insulin plasma levels over time (*G* and *I*), glucose-dependent insulin secretion (*Y*), and total insulin secretion (*S*). While the ODEs are deterministic, their free parameter values are uncertain: they were obtained by fitting noisy and sparse measurements of insulin and glucose levels (20); and they do not account for variability in insulin response as a function of hidden (unseen) variables, such as an individual, time of day, and meal composition other than glucose. To reflect the uncertainty in the free parameters, we specified a prior distribution over each parameter value. In addition, we used a DBN to describe the change in insulin and glucose levels over time, given these parameters and glucose intake during a meal. Lastly, we introduce a boolean variable *T2D*, indicating a diabetic or healthy subject. Thus, the surrogate model now accounts for both the large uncertainty in the model parameters and the statistical dependencies among the model variables over time (Fig. 2).

### Step 2: Coupling surrogate models

Surrogate models enable us to couple multiple input models through subsets of statistically related variables. Their coupling requires some shared reference variables (*i*.*e*., coupling variables). Suitable coupling variables can often be found with the aid of a high-resolution representation of the physical world (*e*.*g*., atomic coordinates in space and time) or some function of these variables (*e*.*g*., coarse-grained coordinates over particles or time). These latent (hidden) variables serve only to formally relate variables from different surrogate models; their values do not need to be known. To couple variables from two or more surrogate models, we describe their relations with the coupling variables, as follows. First, we identify subsets of potentially related variables from multiple surrogate models, as currently determined by an expert based on prior knowledge. Second, for each such subset of surrogate variables, we define corresponding coupling variables. Finally, we devise conditional PDFs (couplers) on each subset of surrogate and coupling variables. We aim to couple as many surrogate models with each other as possible, culminating in a joint PDF over all surrogate models. Importantly, there is generally not one correct choice for the coupling step. Instead, coupling is an external modeling choice and a model in and of itself, just like the input and surrogate models (*M*_*C*_ in Fig. 2). Automated methods for performing this step are conceivable (Discussion). In addition to priors corresponding to input models, we also use data likelihoods when convenient to define couplers (*e*.*g*., GI data model, Fig. 2).

### Example

Four of the eight surrogate models in the metamodel include variables referring to rates of insulin secretion, although in different contexts and spatiotemporal scales. The postprandial response (PR) and pancreas (Pa) surrogate models include a variable referring to the total secretion rate from pancreas to plasma (*S*^*PR*^ *and* 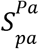, respectively; Fig. 2). The pancreas (Pa), vesicle exocytosis (VE), and GSIS signaling (Sg) models include a variable referring to the secretion rate from a single β-cell (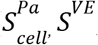, *and S*^*Sg*^, respectively). To relate these variables, we introduce two coupling variables: the true insulin secretion rate from pancreas to the portal vein averaged over population and time 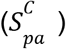 and the true (but unknown) secretion rate from a primary pancreatic β-cell to the extracellular matrix averaged over population of cells 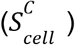. Finally, we impose conditional PDFs on subsets of these surrogate and coupling variables, relying on the pancreas model as a straightforward bridge between different scales. At the plasma level, *S*^*PR*^ in the pancreas response model is conditionally dependent on the coupling variable 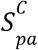, which is in turn conditionally dependent on 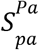 in the pancreas model. At the cell level, 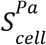 is conditionally dependent on 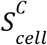, which is in turn conditionally dependent on *S*^*VE*^ and *S*^*Sg*^. Thus, four surrogate models, each describing different scales and aspects of insulin secretion, are coupled. This example provides a blueprint for how more complex models of variation among cells and individuals can also be included. For example, reference variables may describe secretion rates for different individual cells within a single islet or different cell lines.

### Step 3: Harmonize input models by backpropagation of updated variable PDFs

In the last step, information in the coupled surrogate models is propagated back to the original input models. This update is achieved by first updating surrogate models (Fig. 2). A surrogate model PDF can be updated by either marginalizing out or conditioning on each variable in all other surrogate models. In fact, a PDF spanned by any combination of variables from any surrogate models can be computed by marginalizing out and/or conditioning on the other surrogate variables. Finally, we update each input model by relying on a mapping between the surrogate and input model variables. Alternative backpropagation schemes can be performed in parallel (*e*.*g*., conditioning on different values of some variable). Other backpropagation schemes may be explored in the future (Discussion).

### Examples

The postprandial response model includes an input parameter for basal plasma glucose level,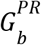 (Fig. 2) (20). Its surrogate model includes a corresponding variable that is distributed normally (mean value of 5.1±1 mM and 9.2±1 mM for healthy and diabetic individuals, respectively), thus describing its prior uncertainty. Following the coupling step, we obtain a joint PDF spanning variables in all surrogate models, including 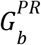. To update a 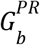 estimate for the postprandial response surrogate model, we compute its marginal PDF from this joint PDF, conditioned on the variable indicating a healthy or diabetic individual. The *G*_*b*_ parameter in the original postprandial response model is then replaced with the mean estimate of 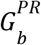 in the surrogate model. This process is repeated for other free parameters of the postprandial response model individually or jointly. Either way, as a result, the updated postprandial response model reflects information from all other input models, *via* the coupling of insulin secretion rates in different surrogate models performed in Step 2.

Another example is provided by the vesicle exocytosis model, which specifies positions of thousands of cellular components over millions of Brownian dynamics trajectory frames (Step 1). As discussed above, the surrogate model has significantly fewer variables. Nonetheless, the PDF of the surrogate model itself encodes the statistical relations among key free parameters and other variables of the input model (example in Step 1). Thus, useful information can be extracted directly from the PDF of the harmonized surrogate model. For example, an updated estimate of vesicle trafficking rate in the vesicle exocytosis model 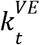 as a function of insulin secretion rate over time in the postprandial response model *S*^*PR*^ can be computed directly from the marginal PDF of 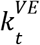, conditioned on *S*^*PR*^ (Fig. 2, backpropagation step). In addition, estimates of the corresponding free parameter *k*_*t*_ and other free parameters of the original input model are updated by backpropagation from the harmonized surrogate model, followed by recomputing spatiotemporal trajectories of vesicle exocytosis using these parameters, now harmonized based on all other input models.

### A proof-of-concept Bayesian metamodel of GSIS

By applying conversion, coupling, and backpropagation (Fig. 2), we divide-and-conquered the task of modeling GSIS, thus decentralizing required computing and expertise. We now discuss how Bayesian metamodeling produces a more complete description of GSIS than the original input models; contextualizes them; increases their accuracy and precision; and resolves conflicting information in the input models. Several simplifying assumptions in the input, surrogate, and coupling models are deemed acceptable at the present time, because the current purpose is to illustrate metamodeling rather than advance knowledge about GSIS.

### Completeness and contextualization

Completeness of a model is the degree to which the model describes all relevant aspects of the modeled system, given the questions asked. By construction, Bayesian metamodeling provides a more complete description of GSIS than any of the input models on their own. It also contextualizes the different input models by relating previously uncoupled variables (from different input models) to each other. As a consequence, Bayesian metamodeling can be used to assess the effect of different models on one another, and augment each model with information in other models to which it was oblivious prior to metamodeling.

### Examples

Incretins, such as the GLP1 peptide (21), are hormones secreted from the endocrine pancreas that regulate plasma glucose levels (22). In the presence of glucose stimulus, GLP1 increases insulin secretion by activating GLP1R, the cognate receptor of GLP1 on pancreatic ß-cells. Indeed, GLP1R agonists are commonly used to treat type 2 diabetes (21), although clinical significance of this activation in ß-cells *versus* other tissues is yet to be determined (23). The postprandial model (20) does not include any information on GLP1, GLP1R, nor their effect on systemic insulin response after a meal (Fig. 2; SI Appendix: Supplementary Text 1.1). In contrast, this information is included in the GSIS signaling model, which describes how GLP1 activates GLP1R, insulin biosynthesis, and secretion pathways downstream of GLP1R, within a single ß-cell (Fig. 2; SI Appendix: Supplementary Text 1.4). In the metamodel, variables from both models are coupled (Fig. 2). This coupling enables us to re-estimate the free parameters of the postprandial model for different choices of extracellular concentrations of GLP1 (Fig. 3B). Consequently, the updated postprandial model successfully recapitulates the incretin effect (*i*.*e*., the empirically observed effect of elevated GLP1 concentrations on postprandial insulin and glucose levels) (21); in other words, the postprandial model was contextualized by the model of GSIS signaling.

**Figure 3.**
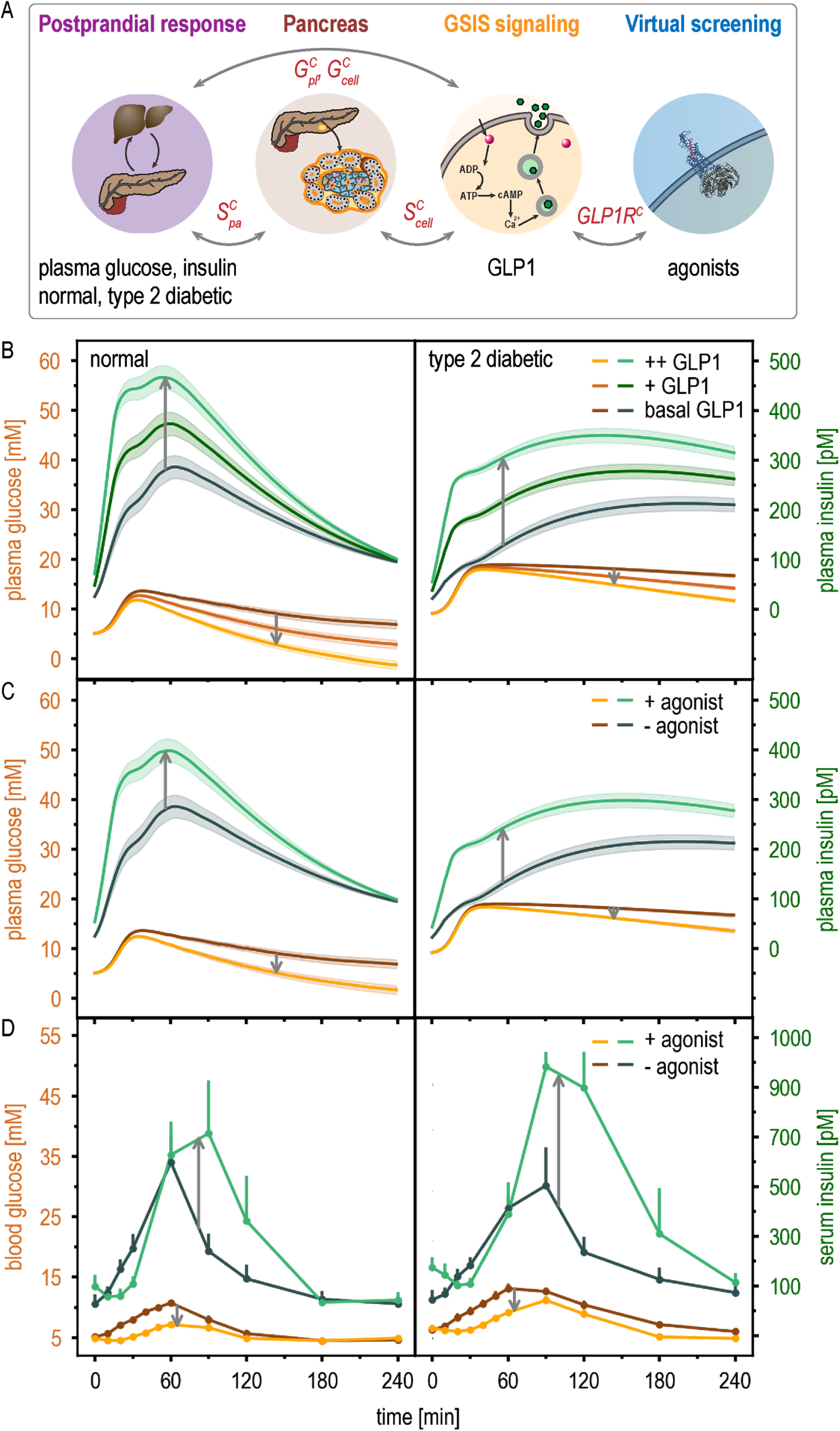
Contextualization of input models by metamodeling is illustrated by the effect of GLP1 and incretins on GLP1R. (A) Coupling among four input models is indicated schematically. Gray arrows indicate the flow of information between the models, *via* the coupling variables in red. Time courses of postprandial glucose (orange shades) and insulin (green shades) plasma levels are shown for normal (left) and type 2 diabetic subjects (right). (B) Metamodeled time courses are shown for three glucagon-like peptide 1 (GLP1) concentrations in the GLP1 data model: basal, medium (+), and high (++). The shaded areas indicate standard deviation in the posterior PDFs. (C) Metamodeled time courses are shown for postprandial response with and without a GLP1R agonist in the virtual screening model, using analogue M2 in the virtual screening library (93). (D) Experimental time courses are shown for postprandial response with and without a GLP1R agonist, exenatide (synthetic Exendin-4) (23).

A second example is provided by the GLP1R model, which is an atomic spatial model of GLP1R activation by GLP1 analogs (Table 1). In the GSIS metamodel, this input model augments both the postprandial and the GSIS signaling network models with binding affinities of various GLP1 analogs for GLP1R, as predicted by virtual ligand screening (Fig. 5A; Table 1). The GLP1R model thus facilitates predicting the effect of GLP1 analogs on insulin secretion at the systemic and cellular levels, respectively (Fig. 3C). These predictions recapitulate the clinical observations of the effect of GLP1 analogs on postprandial insulin and glucose levels (Fig. 3D). This example also illustrates the modularity of metamodeling: Additional input models can be incorporated into an existing metamodel, iteratively increasing its completeness.

### Effect of metamodeling on accuracy and precision

A useful model needs to be sufficiently accurate and precise, given the questions asked. Metamodeling aims to increase the accuracy (decrease the systematic error) of variable estimates as much as possible, given the accuracy and precision of the input models.

Accuracy and precision of metamodeling can be benchmarked in two ways, as is the case for any modeling method. First, a metamodel can be validated retrospectively by comparison against an independently determined reference (*e*.*g*., the validation of the GSIS metamodel by experiment in Fig. 3D). Second, the accuracy of metamodeling can be assessed with a synthetic benchmark. In such a benchmark, true values of free parameters for the various input models are defined, followed by enumerating the input models and the corresponding output metamodels for combinations of input free parameter values. We can then systematically assess the impact of metamodeling on the accuracy and precision simply by comparing the output joint PDFs in the corresponding metamodels with the true values of the free parameters. Consistently with the broad definition above, systematic error of a variable is defined specifically as the difference between the mean of its output PDF and the true value, indicated by 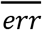 and precision (random error) is defined as the standard deviation of its output PDF, indicated by *σ*.

### Example

In a synthetic benchmark, we assess the impact of metamodeling on the systematic and random errors of free parameters 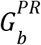 in the postprandial model and 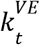 in the vesicle exocytosis model. 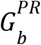 corresponds to the basal glucose level in plasma; and 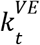 corresponds to the effective rate of vesicle trafficking towards the cellular periphery. Prior uncertainties in their values (*e*.*g*., due to variation among individuals and over time) are reflected in their input PDFs; for example, the PDF for 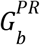 of healthy individuals is a Gaussian distribution with the mean of 5.1 mM and the standard deviation (*σ*) of 1.0 mM (SI Appendix: Table S1). In the metamodel of GSIS, 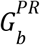 and 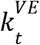 are coupled indirectly *via* reference variables (Fig. 2). As a result of metamodeling, the systematic and random errors of both variables may in principle either increase, decrease, or remain constant, depending on the magnitude and directionality of the errors in the prior estimates of 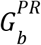 and 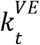 (Fig. 4; SI Appendix: Fig. S14). To illustrate this general point, we compute actual changes in the systematic and random errors of 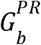 and 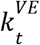 produced by GSIS metamodeling (output accuracy and precision), as a function of the accuracy and precision of 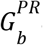 and 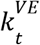 in the input models (input accuracy and precision).

**Figure 4.**
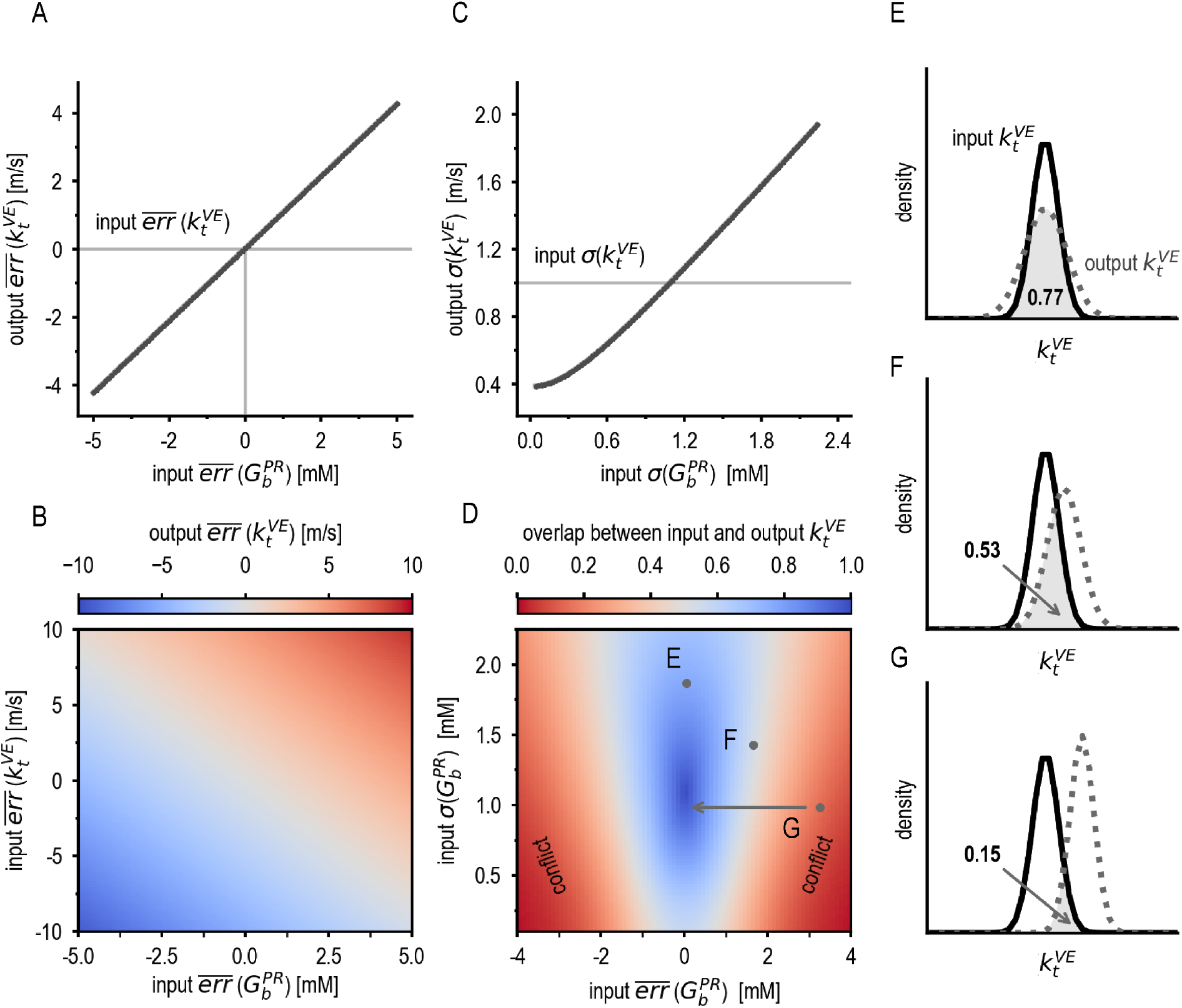
Effect of metamodeling on model accuracy (systematic error) and precision (random error). (A) Statistical dependency of the output systematic error 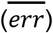 of the variable 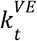 in the vesicle exocytosis model on the input systematic error of the variable 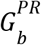 in the postprandial response model. The coupling coefficient corresponds to the slope of the line. (B) The output systematic error of 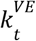 given different input systematic errors of 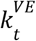 and 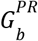. (C) Statistical dependency of the output random error (*σ*) of 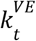 on the input random error of 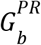. (D) The overlap between input and output 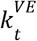, as a function of input systematic error (x-axis) and random error (y-axis) of 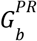. Conflicting models correspond to the red areas. (E, F, G) The input and output PDFs of 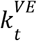 corresponding to points E, F, and G in (D), respectively. Arrow in (D) indicates the direction of resolving conflict by improving the accuracy in input 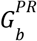. All output values are at *t* = 100 min.

We first discuss the output accuracy (systematic error) as a function of the input accuracy. A coupling coefficient of a variable with respect to another variable is defined as the sensitivity of systematic error in its output PDF to systematic error in the input PDF of the other variable (slope in Fig. 4A; SI Appendix: Fig. S14A). As expected, the magnitude of the coupling coefficients of 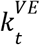 with respect to 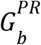 and 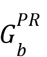 with respect to 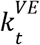 is relatively high (0.84 m/s per mM and 0.25 mM per 1.00 m/s, respectively). Indeed, slower trafficking of insulin granules may lower insulin secretion rate in dysfunctional ß-cells (24), potentially explaining elevated basal glucose levels in the plasma of diabetic individuals. Thus, metamodeling correctly couples two variables that were not coupled before metamodeling (because they occured in separate input models). The coupling coefficient of 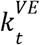 with respect to 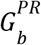 is positive, because these two variables are positively correlated in the metamodel. Thus, when 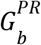 is overestimated and 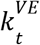 is underestimated in their input models, or *vice versa*, these two estimation errors are likely to diminish each other in metamodeling (Fig. 4C, gray diagonal). Conversely, when both are either underestimated or overestimated, metamodeling likely decreases their accuracy (Fig. 4B, red and blue regions). Nonetheless, in analogy with the law of large numbers (25), we expect that the larger the number of input models, the more likely the random errors in the input models cancel out, in turn leading to more accurate estimates in the metamodel.

Next, we discuss the output precision as a function of input precision. When the random error of 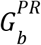 in the input model (input σ) increases, the random error of 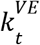 in the output metamodel (output *σ*) also increases (Fig. 4C). Thus, input models with lower random error contribute to lower random error of variables from other models. In other words, metamodeling correctly weighs the uncertainties of the different input models, and updates output precisions accordingly. In contrast, when the input random error of 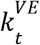 increases, the output random error of 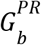 still increases, but significantly more slowly (SI Appendix: Fig. S14). A possible explanation is that 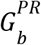 is stabilized through its coupling to variables from multiple models (*e*.*g*., the GSIS signaling model and pancreas model), and is thus less sensitive to random errors in coupled variables from a single input model. This observation illustrates one potential benefit of weighing information from multiple models *via* metamodeling.

### Conflicting models

Changes in variable estimates after metamodeling can be used to identify conflicts between a variable in one input model and other input models. After metamodeling, a variable PDF may change significantly relative to its precision (Fig. 4D-G). Thus, conflict between the variable and other input models is quantified by the overlap between variable PDFs before and after metamodeling (Fig 4D) (26). When the systematic error of 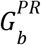 in the input postprandial model is low (point E in Fig. 4D), it is consistent with the value of 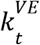 in the input vesicle exocytosis model; consequently, the overlap between its input and output PDFs is high (Fig. 4E). Even when the input systematic error of 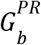 is high, but its input random error (*σ*) is also high (point F in Fig. 4D), the overlap between the input and output PDFs for 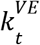 remains relatively large, indicating conflicting information that is tolerable given the high prior uncertainty in 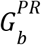 (Fig. 4F). In contrast, when the input systematic error of 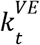 is high and its input random error is low (point G in Fig. 4D), the overlap between the input and output PDFs for 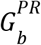 becomes smaller, indicating conflicting information that is not tolerable given prior uncertainty in 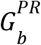 (Fig. 4G). Thus, tolerability of conflicting information in different input models is identified by comparing the overlap among PDFs before and after metamodeling. Moreover, variables leading to untolerable conflicts can be prioritized for experimental followup to refine the input models, and thus resolve the conflicts. For instance, given a conflict between 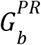 and 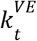, an improved measurement of 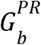 could result in a refined postprandial model with better accuracy and precision, removing the conflict (arrow from point G in Fig. 4D). Lastly, as with accuracy, introduction of additional models to the metamodel can also resolve such conflicts by providing an additional source of information about conflicting variables.

## Discussion

### Summary

Here, we developed Bayesian metamodeling, a divide-and-conquer approach to modeling complex systems, such as the cell. Metamodeling is not meant to replace other modeling methods, including cell modeling methods. Instead, it is meant to integrate, refine, and harmonize all other relevant models. Next, we discuss (i) combining different models, (ii) the relationship of metamodeling with other whole-cell modeling approaches and integrative modeling, (iii) the advantages of metamodeling, (iv) major limitations of metamodeling and how they might be addressed, and (v) the application of metamodeling to cell modeling by the Pancreatic ß-Cell Consortium.

### Combining multiple models

Combining multiple models using the same representation (*i*.*e*., same type of model, same modeled system) is performed relatively often with the goal of increasing the accuracy and precision or the coverage of the combined model. For example, different protein structure models can be averaged into a hopefully more accurate and precise average model (27); multiple types of classification models can be combined to obtain a more accurate classification model (28); multiple cellular networks increase the coverage of the cell (8); and docking of multiple subunit structures results into a model of the complex (29). Likewise, multiple models of different types can also be combined to get a model that describes a larger system and describes it more comprehensively. For example, the 2013 Nobel prize in chemistry was awarded to M. Karplus, M. Levitt, and A. Warshel for harmonizing quantum mechanics and molecular mechanics, thus providing an early example of coupling and multiscaling at atomic resolution (2, 3); and different types of models are combined for weather prediction (30). In an early example of multiscale modeling of GSIS, first crystallographic structures of insulin and glucagon gave rise to more holistic, functional depictions of signalling and storage in insulin granules (31). Cell mapping in particular has also been addressed by combining models of different representations. A groundbreaking method propagates a complete cell model from an initial time point by using output from some models as input for other models at the next time point on the trajectory, with different models being coupled *via* metabolites (9, 11). In a second example, stochastic reaction diffusion master equation (RDME) models of chemical reaction networks, describing the formation of splicing machinery, were combined with a spatial model of the HeLa cell to study the influence of spatial organization on splicing, based on data from cryo-electron tomography, mass spectrometry, fluorescence microscopy / live-cell imaging, and -omics (10). However, to the best of our knowledge, a general approach to combining heterogeneous input models of any type or scale into a unified model does not yet exist.

Here, we formalized the model integration problem in general terms and described one practical approximate solution, termed Bayesian metamodeling. The solution depends on the universality of representing the statistical uncertainties and dependencies among the variables spanning any model or dataset. As a result, any model or dataset can in principle be input for metamodeling.

### Relationship to other cell modeling approaches

Most generally, any modeling can be seen as sampling of instances of a model of a certain type, using some sampling scheme guided by some scoring function informed by input experimental data and/or prior models. For example, in addition to the above-mentioned approaches, a variety of methods have been used to model various aspects of the cell, based on a variety of data (32–34). Deep-learning approaches were applied to learn cell phenotypes from their genotypes, using a network that mirrors the structural and functional hierarchy of a cell (35), based on genomics and proteomics data (36). Manual curation was used to construct a repository of genome-scale metabolic models (8), based on various genomics, proteomics, and metabolomics data. A stochastic simulation algorithm was combined with flux balance analysis to model stochastic dynamics of metabolism in *Mycoplasma pneumoniae*, based on metabolic and proteomic data (37). More generally, several modeling platforms were developed for spatiotemporal simulations of reactions, mass transport, and other processes in the entire cell (38–42). Packing algorithms were used to assemble macromolecules in a complete HIV-1 virus particle and *Mycoplasma mycoides* cell at 10-100 nm resolution (43), based on data from structural biology and systems biology. Atomistic molecular dynamics and coarse-grained Brownian dynamics simulations were used to model crowded cytoplasmic environments, resulting in trajectories of millions of particles over microseconds for sections of *Mycoplasma genitalium* (44) and *Escherichia coli* (6). Satisfaction of spatial restraints resulted in architectures of genomes in various types of cells, based on genome-wide mapping of chromatin interactions (45). Similarly, spatial restraints were satisfied to create a snapshot of a synaptic bouton at atomic resolution based on data from quantitative immunoblotting, mass spectrometry, electron microscopy, and super-resolution fluorescence imaging (46). A number of methods rely on image processing or machine learning from images. For example, 3D reconstruction and segmentation were used to create a model of mouse pancreatic ß-cell ultrastructures using data from serial section electron tomography (47); convolutional neural networks were applied to compute fluorescent 3D cellular maps from 3D label-free transmitted-light live-cell images or 2D electron microscopy images (48); image processing and machine learning techniques were used to compute subcellular sarcomeric organization states in cardiomyocytes based on data from single-cell RNA sequencing and quantitative imaging of gene expression, transcript localization, and cellular organization (49); finally, a pipeline for multiplexing different imaging modalities was used to map protein-ultrastructure relationships from cryogenic super-resolution fluorescence microscopy and focused ion beam–milling scanning electron microscopy (50). Thus, most cell mapping approaches are limited in the types of cell representation and reliance on limited types of data and/or prior models. In contrast, metamodeling can in principle use any set of representations that can be informed by the available data and are useful for addressing given biological questions.

### Relationship to integrative modeling

As mentioned in Introduction, metamodeling is a special case of integrative modeling. In addition to integrative structure modeling (4), other variants of integrative modeling include integrative pathway mapping (51), modeling of spatial organization of genomes (52), integration of imaging and -omics data (53), studying of splicing codes based on multiple sources of data (54), integration of single cell transcriptomic, epigenetic data, and protein counts (55), integration of multimodal neuroimaging data (56), and general machine learning techniques for dealing with multimodal data (18). Bayesian metamodeling is in fact a decentralized form of integrative modeling in which the focus is shifted from integrating data to integrating prior models. In addition to using data to compute input models, data can also be used as an input model itself, as exemplified by the GI data and GLP1 data models (Fig. 2; Table 1; SI Appendix: Supplementary Text 1.7 and 1.8); in other words, data can be incorporated directly *via* data likelihoods in the joint PDF during the coupling stage. Thus, an integrative modeling problem can also be formulated as a metamodeling problem, benefitting from the advantages of its divide-and-conquer strategy.

### Advantages of metamodeling

We outline here a number of advantages of metamodeling over more centralized approaches to data integration: First, metamodeling is highly modular and benefits from heterogeneity of representations. Different aspects of the cell and its functions are modeled by different methods, informed by different data, and represented with different variables at varying levels of granularity (Figs. 1 and 2). Second, metamodeling facilitates multiscaling. This advantage arises from modularity that also allows combining models at different scales. Third, metamodeling is computationally efficient. The large task of computing a model of the cell is distributed among smaller parallel computations required to compute individual input models. Fourth, metamodeling is collaborative. It allows autonomous contributions by different research groups with expertise spanning diverse scientific disciplines, thus maximizing flexibility, scalability, and efficiency among collaborating experimentalists and modellers. Fifth, metamodeling is statistically objective. This objectivity derives from the use of Bayesian formalism for modeling the relations among different system parts. Sixth, metamodeling increases model completeness. A metamodel provides a maximally complete view of all cellular aspects, given the available input models (Figs. 3 and 4). Seventh, metamodeling can couple previously independently modelled cellular aspects. This advantage results from harmonizing different input models with respect to each other, thus providing more context for each input model (Fig. 3). Eighth, metamodeling often improves accuracy and precision. This improvement is achieved by updating model variables and their uncertainties by considering information from multiple input models, thus often improving the final estimates (Fig. 4A-C). Finally, metamodeling helps with resolving conflicts among input models. If different input models contain contradictory information, metamodeling highlights these inconsistencies and thus helps identify new experiments that may resolve them (Fig. 4D). Next, we discuss a few of these advantages in more detail.

### Modularity

Bayesian metamodeling can in principle use any type of an input model, including deterministic or stochastic, static or dynamic, and spatial or non-spatial. The only requirement is that an input model is specified quantitatively. Importantly, metamodeling does not require the data used to construct each model. Therefore, input for metamodeling can be obtained relatively easily from independent research groups that do not necessarily collaborate or share expertise. Moreover, a metamodel can be updated iteratively with additional models, utilizing new datasets, technologies, and modeling techniques as they emerge. Thus, metamodeling enables a plug-and-play approach to building complex models from simpler models.

To illustrate the benefits of this modularity, we now discuss practical examples of upgrading the current GSIS model to better account for variation across (i) multiple cells of the same type and (ii) different types of the cell. Each one of these variations can be modeled by either improving an existing input model or by adding a new input model, without changing other input models, as follows. First, variation across multiple cells of the same type could be modeled by replacing the current linear pancreas model with a more elaborate model that accounts for electrical synchronization in networks of ß-cells (57) and data on the role of leader ß-cells in these networks (58); such a model could be coupled to insulin vesicle exocytosis and/or GSIS signaling models, each parametrized to reflect cell variation. Thus, metamodeling may provide an effective path to investigate the source of cell heterogeneity in glucose responsiveness, an open question of great biological interest (59). Second, variation across cell types could be accounted for by including a separate input model for each type of the cell (*e*.*g*., primary ß-cells and insulin-secreting model cell lines, such as INS-1 and INS-1E tumor cells (60, 61)). During the coupling stage, the weight of variables from each input model should reflect the confidence in it. For instance, data from model cell lines is often considered significantly less informative about primary cells than the data from the primary cells themselves (62). Indeed, variables from the insulin metabolism model, which was informed by experiments in INS-1E cells, are only weakly coupled to variables from other models (SI Appendix: Tables S10-S11). Other variations, such as those among different individuals, can in principle also be addressed similarly.

### Multiscaling

Metamodeling can couple different input models despite significant differences in their scales. In fact, even the current GSIS metamodel covers the scales from atomic and femtoseconds of molecular dynamics simulations to the whole body and hours of the postprandial response model (Table 1). Thus, multiscaling is another advantage of modularity. This coupling is achieved by imposing statistical correlations among variables on different scales. Thus, metamodeling may bypass the need to compute the propagation of signals across scales explicitly, which typically necessitates specialized model representations and algorithms (63). For example, the pancreas model provides a simple description of the expected statistical relation between secretion of insulin at the cell and systemic levels, thus helping to couple the postprandial response and GSIS signaling models. Likewise, the GLP1R model at atomic scale is coupled to all other models *via* the GSIS signaling model at cellular scale by imposing expected statistical correlation between receptor activation by a small molecule agonist and activation of GSIS signaling in the cell. This coupling allows us to use the GSIS metamodel to correctly predict the effect of incretins and other small molecule ligands on systemic insulin response (Fig. 3).

### Facilitating community collaboration

Due to its modularity, metamodeling is expected to provide a community tool for contributing to whole-cell modeling, as exemplified by its use within the Pancreatic ß-Cell Consortium (14). We developed tutorials serving as onboarding material to allow others to contribute their input models (SI Appendix: Supplementary Text). In the future, we will also create a website to serve as a graphical user interface and develop methods for automated conversion of input models into surrogate models. This functionality will provide non-experts in computational modeling with an opportunity to contribute and improve their individual models. At its core, metamodeling is rooted in collaboration and appreciation for the details of disparate data, methods, and models, which cannot be achieved by any individual scientist, research group, or institution. To further support the collaboration, the Pancreatic ß-Cell Consortium is creating cyberinfrastructure for archiving and disseminating experimental data and models that will be integrated with metamodeling. Thus, each time an input model, surrogate model, and/or a coupler is provided or upgraded, the metamodel and input models can be updated automatically.

Many of the most important questions in biology are centered around issues of data integration and at the intersection of multiple fields. Thus, the development of methods and tools that build bridges between siloed research is essential. An existing example is the Protein Data Bank (PDB) of known protein structures, which in many ways nucleated the structural biology community (64). Indeed, the latest effort of the PDB to support integrative structures based on varied data from multiple methods (65) is narrowing the gap between the PDB and the whole-cell mapping. Other key community resources provide for standardization, archival, and dissemination of models, thus facilitating explicit and implicit collaboration among a diverse set of researchers (8, 12, 66–68).

### Limitations of metamodeling

While Bayesian metamodeling can in principle be used to couple any set of input models, it is not always clear that they can be coupled usefully. Next, we identify a number of limitations of the current implementation of metamodeling and outline how they might be addressed.

First, to incorporate complex input models more accurately, alternative approaches for converting these models into a unified surrogate probabilistic representation should be explored (Step 1). While nonlinear models can be approximated by DBNs (Fig. 3), other methods for learning complex PDFs spanned by a large number of inter-dependent variables include a non-linear implementation of PGMs (github.com/tanmoy7989/bayesian_metamodeling_tutorial). In addition, deep-learning approaches, such as variational autoencoders, generative adversarial networks, and temporal variants might also be useful, although they generally require a large amount of training data and they are not always easily interpretable (18). Nonetheless, deep neural networks have already been shown to provide practical solutions for representing low-dimensional surrogate models for complex physical systems (69). Finally, non-probabilistic approaches, such as integer programming (70), might also be explored.

Second, only a limited set of coupling schemes have been used so far, based on imposing statistical dependencies *via* PGMs (Step 2). While PGMs and other probabilistic approaches (*e*.*g*., generative deep learning models) provide a relatively general solution for coupling models of any type, some types of models may be coupled more efficiently and/or accurately by other types of couplers. For example, the coupling of ligand and receptor structural models in molecular docking can be achieved naturally, accurately, and efficiently *via* minimizing the free energy of the complex (71). Thus, future work should explore additional types of couplers for common types of models. As a special case, couplers for input models at different spatiotemporal scales should be improved. Multiscale integration is currently performed *ad hoc*, based on prior knowledge about expected correlations across scales. Standardized schemes and automated methods for integrating models across scales should be developed, including more efficient representations for multiscale PDFs. As discussed above, at the very least, metamodeling facilitates a formal integration by imposition of statistical correlations across scales in cases where explicit physically-inspired coupling is not yet possible.

Third, to maximize modeling accuracy and precision, metamodeling should be guided by formal optimality criteria (loss or fitness functions) for (i) ranking surrogate models, (ii) reference variables used to couple the surrogate models, (iii) the couplers themselves, and (iv) the backpropagation scheme. For example, good surrogate models should recapitulate statistical dependencies in the input models, and good couplers may be required to recapitulate experimental data on statistical dependencies among input models.

Fourth, the metamodeling process should be entirely automated, in part benefiting from the optimality criteria above. Such automation will require sampling in the space of alternative metamodels to choose an optimal metamodel, for example, by sampling the free parameters and topologies of surrogate models and couplers (17). Automation will also be facilitated by developing tools that streamline interactions with non-experts in modeling (Facilitating community collaboration, above).

Fifth, we should develop methods to validate a metamodel. Although the uncertainty of a metamodel is already quantified by its PDF, additional assessment may be useful. A relatively general approach to model assessment has been developed for integrative structure modeling, quantifying the degree of sampling exhaustiveness, the match between the model and the data used to construct the model, the match between the model and the data not used to construct the model, and model uncertainty (72). In metamodeling, additional opportunities for assessment include identifying conflicts between input models and assessing error propagation (Fig. 4). For example, while our results indicate that random or uncorrelated systematic errors in different models are likely to be averaged out through their coupling, such coupling may also lead to amplification of error in one input model as it propagates across models. Methods for detecting and minimizing such errors should be developed, possibly by borrowing from methods for stabilization of dynamic systems (73).

Finally, our primary purpose here was to illustrate metamodeling rather than advance our understanding of GSIS biology. Thus, our current metamodeling relies on a small set of relatively simple input models and numerous simplifying assumptions, some of which are summarized in SI Appendix. While even this simplified metamodel has been validated by data not used in its construction (Fig. 4), we have not yet obtained any new insights into GSIS. Future implementations of metamodeling should be tested using a larger number of input models of higher complexity.

### Future application to whole-cell modeling

Together with the entire Pancreatic ß-Cell Consortium (14), we are working to enrich the current GSIS metamodel with additional input models based on diverse types of data to create a more accurate, precise, and complete model of the pancreatic ß-cell. These additional input models cover key aspects of GSIS biology in health and disease, including glucose sensing (74, 75); insulin vesicle biosynthesis, trafficking, docking, and exocytosis (76); recycling of misfolded proteins by proteasomes and autophages (77); membrane phospholipid biosynthesis at mitochondria-associated endoplasmic reticulum membranes (MAMs) (78); regulation of intracellular calcium flux from ER to mitochondria (79); global spatiotemporal dynamics of islet insulin secretion (80); pulsatile insulin secretion (81); interaction with hepatocytes (82); phosphoproteome map (83); and spatial genome organization (84). The upgraded metamodel is expected to be useful for designing more effective future experiments, discovering biological mechanisms, and generating hypotheses, which will in turn enhance the model itself. We also anticipate metamodeling of ß-cells by the Pancreatic ß-Cell Consortium will serve as a template for modeling other types of cells and, indeed, other complex systems.

## Methods

The software, input files, and example output files for the present work are available at github.com/salilab/metamodeling. The metamodel was implemented using the BNET package in MATLAB by Kevin Murphy, github.com/bayesnet/bnt (commit 21dfdfa) with minor modifications of the DBN module (github.com/salilab/metamodeling/tree/master/bnt-master); the probabilistic graphical models in the tutorial (github.com/tanmoy7989/bayesian_metamodeling_tutorial) were implemented in the Python package PyMC3 (version 3.8) (github.com/pymc-devs/pymc3/releases/tag/v3.8). For an outline of the approach, see Results; for details, see SI Appendix.

## Acknowledgments

We are grateful to all members of the Pancreatic ß-Cell Consortium for providing the context in which this research was performed. In particular, we appreciated the discussions with Helen Berman and Peter Butler. We also acknowledge helpful comments by Keren Lasker, Trey Ideker, Marcus Covert, Eran Agmon, Reshef Mintz, and Thomas L. Blundell. The work was funded by grants NIH/NIGMS R01GM083960, NIH/NIGMS P41GM109824, and NIH/NIAID U19AI135990 (AS), Bridge Institute at USC (AS, RCS, and KLW), ShanghaiTech University (LS, CW, JZ, and AL), Bridge Institute postdoctoral fellowship (KP), the Burroughs Wellcome Fund Travel Award (KLW), and a starting grant from the Hebrew University of Jerusalem (BR). We also acknowledge the High Performance Computing (HPC) Platform of ShanghaiTech University for computing time.

## Author Disclosures

RCS acknowledges that USC is his primary academic affiliation.

## SI Appendix

### Supplementary Text

We first describe each of the eight input models and their corresponding surrogate models (Section 1), followed by outlining how surrogate models are coupled (Section 2).

#### 1. Input and surrogate models

##### Input models

Our current goal is to illustrate metamodeling with a set of input models, without necessarily improving our understanding of pancreatic β-cell biology. Thus, we do not discuss in detail the validity of previously published input models (20)(87)(20). We do, however, highlight the main assumptions in constructing the previously unpublished input models (the pancreas, GSIS signaling, insulin metabolism, virtual screening of GLP1R, GI data, and GLP1 data models).

#### Surrogate models

We outline each surrogate model and how it was constructed from the input model, including modeling assumptions. We also provide a table listing the conditional PDF of its variables at different time slices of its DBN and their parameters.

#### DBNs of surrogate models

As described in the main text, a surrogate model is represented by a PDF over some variables of the corresponding input model and potentially additional auxiliary variables. A surrogate model aims to approximate statistical dependencies among the variables in the original model, potentially in a simplified form. Each surrogate model was represented by a DBN (17). Briefly, a DBN factorizes a PDF over a set of time-dependent variables by describing a conditional PDF for each random variable at time slice *t+Δt* as a function of some random variables at time slice *t* and/or time slice *t+Δt*. In addition, a DBN describes a conditional PDF for all variables at an initial time slice *t*_*0*_. If the values of some variables are observed, posterior conditional or marginal probabilities can be inferred for any subset of variables in the DBN at any time point. In addition, DBN parameters and topology can be learned from observed data.

#### DBN implementation

The topology of the DBN for each surrogate model is given in the top panel of Figure 2. The software implementation, input files, and sample output files are available at github.com/salilab/metamodeling. The conditional PDF of each random variable is a normal distribution whose mean is the weighted sum of some random variables (a linear Gaussian) in the current and/or previous time slice, with manually assigned standard deviations; the use of non-linear models is illustrated in a tutorial (github.com/tanmoy7989/bayesian_metamodeling_tutorial). The discrete time step *Δt* for the DBNs of all surrogate models was set to 1 min, although the original input models are constructed with their own time scales and time granularity. In fact, some models contain time-independent variables, which do not change over the timescale of a model. Time-independent variables may become time-dependent in the coupling stage, due to coupling with time-dependent variables from other surrogate models. To guarantee numerical stability, the conditional PDF of time-independent variables at each time slice is allowed to fluctuate slightly, by assigning them an arbitrary small standard deviation.

### 1.1 Postprandial response model

#### Input model

The postprandial response model describes insulin and glucose levels in the plasma and various body tissues (dependent variables) as a function of time, following a glucose-rich meal, in healthy and T2D subjects (Fig. S1) (20). The values of these variables are computed from the rate of glucose intake (independent variable), using a system of ODEs.

**Figure S1.**
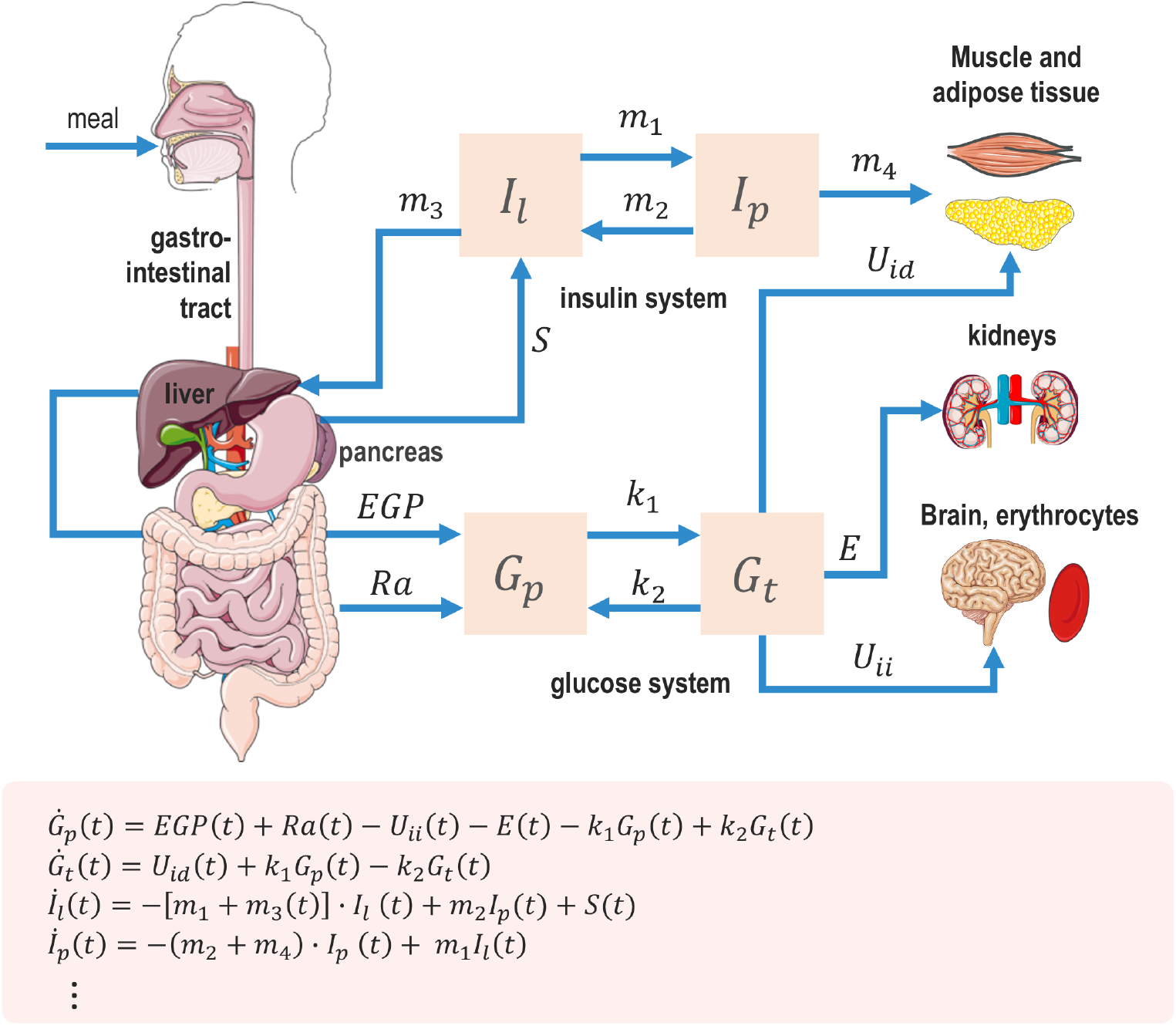
The postprandial response input model. (Top) A schematic of the system of compartments and fluxes, described by a model consisting of 29 ODEs (20). The model takes into account interactions among different physiological systems and organs involved in glucose homeostasis. It was parameterized based on data from a cohort of 204 healthy subjects and 14 T2D subjects. These data include plasma insulin and glucose levels, endogenous glucose production, glucose rate of appearance, glucose utilization, and insulin secretion rate over 420 minutes at 20 minute intervals following a glucose-rich meal. (Bottom) A subset of 4 of the 29 ODEs in the complete model, indicating change in levels of plasma glucose (*G*_*p*_), tissue glucose (*G*_*t*_), liver insulin (*I*_*l*_), and plasma insulin (*I*_*p*_).

#### Surrogate model

The rationale behind the construction of the postprandial response surrogate model is described in Results (second example for Step 1 of metamodeling). It is relatively straightforward to construct a DBN for a system of ODEs, because a DBN can be considered a probabilistic, discretized generalization of a system of ODEs. The postprandial response surrogate model nonetheless simplifies the input model, still capturing key statistical relations among its variables (Fig. 2, Table S1) by fitting some parameters of the conditional PDFs listed in Table 1 to reproduce its dependent parameters. As described above, all conditional PDFs are linear Gaussians.

**Table S1.**
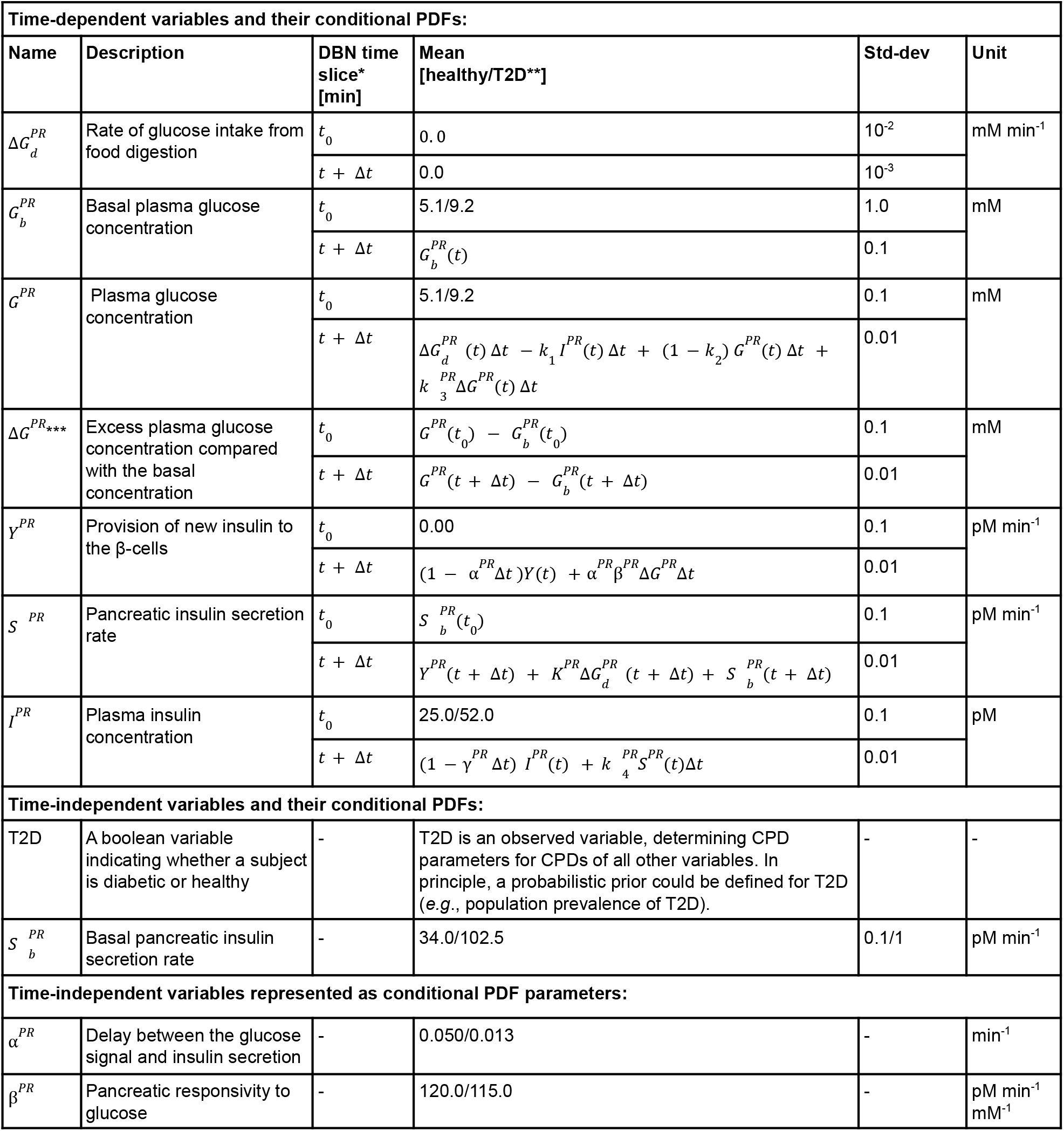
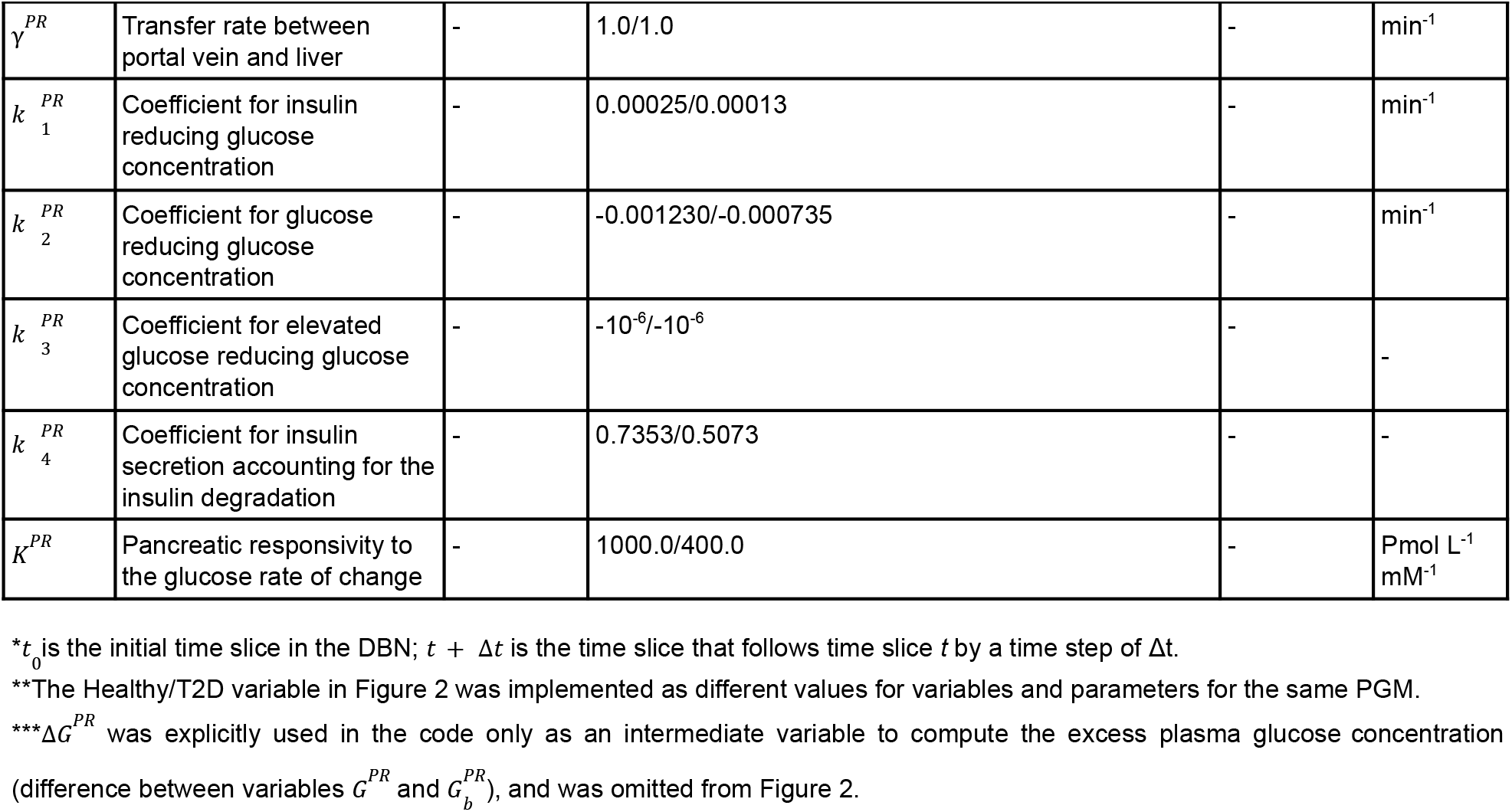
Random variables, corresponding conditional PDFs, and conditional PDF parameters in the postprandial response surrogate model.

### 1.2 Pancreas model

#### Input model

The pancreas model is a simple linear model that relates the insulin secretion rate by individual cells (independent variable) to the insulin secretion rate by individual islets and an entire pancreas (dependent variables) (Fig. S2).

**Figure S2.**
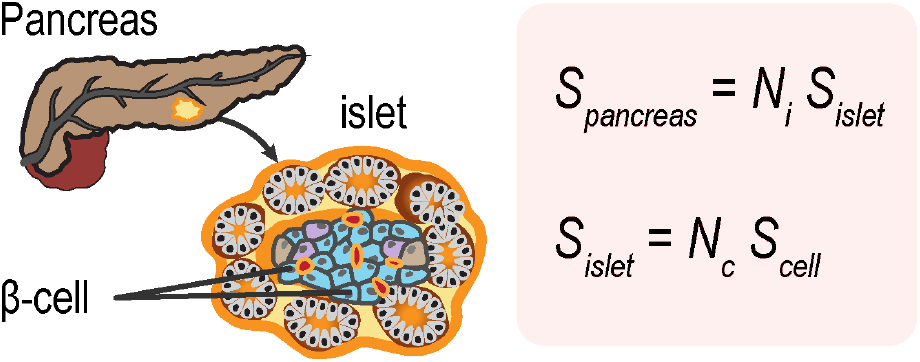
The pancreas input model. The insulin secretion rate of an islet *S* _*islet*_ is the sum of the secretion rates of *N*_*c*_ β-cells (*S* _*cell*_) in the islet; similarly, the insulin secretion rate of the pancreas *S* _*pancreas*_ is the sum of the secretion rates of *N*_*i*_ islets in the pancreas. The model is parameterized based on the estimated *N*_*c*_ =1140 β-cells in an islet (86) and *N*_*i*_ =3.2 million islets in a pancreas (85). Variable descriptions are indicated in Table S2. We made two simplifying assumptions in the construction of the pancreas input model. First, the secretion rates of all β-cells are identical and all islets contain the same number of β-cells. Second, *N* _*c*_ and *N*_*i*_ were assigned identical values for both healthy and T2D subjects, although the proportion of β-cells in T2D islets is marginally decreased compared to normal islets (94).

#### Surrogate model

The DBN describing the pancreas surrogate model is a discretized, probabilistic version of the linear equations of the corresponding input model (Fig. 2, Table S2). It includes conditional PDFs corresponding to linear Gaussians describing the statistical dependencies between insulin secretion rates for β-cells, islets, and the pancreas. This surrogate model is essential for the subsequent coupling of the postprandial response, vesicle exocytosis, GSIS signaling, and GI data surrogate models (Results, Step 2). It includes only time-independent variables. However, it is implemented as a DBN, because its variables become time-dependent during the coupling stage, due to coupling with time-dependent variables of other surrogate models (Fig. 2).

**Table S2.** Random variables, corresponding conditional PDFs, and conditional PDF parameters in the pancreas surrogate model.

| Time-independent variables and their conditional PDFs: |  |  |  |  |
| --- | --- | --- | --- | --- |
| Name | Description | Mean | Std-dev | Unit |
| $S_{cell}^{Pa}$ | Insulin secretion rate of a cell | $S_{cell}^c(t)$ | $10^{-10}$ | pM min<sup>-1</sup> |
| $S_{is}^{Pa}$ | Insulin secretion rate of an islet | $N_c S_{cell}^{Pa}(t)$ | $10^{-6}$ | pM min<sup>-1</sup> |
| $S_{pa}^{Pa}$ | Insulin secretion rate of the pancreas | $N_i S_{is}^{Pa}(t)$ | 0.1 | pM min<sup>-1</sup> |
| Parameters of conditional PDFs: |  |  |  |  |
| $N_c$ | Number of β-cells in an islet | 1140 | - | - |
| $N_i$ | Number of islets in a pancreas | $3.2 \times 10^6$ | - | - |

\*Legend: $t_0$ - the initial time slice in the DBN; $t + \Delta t$ - the time slice that follows time-slice $t$ by a time step of $\Delta t$ .

### 1.3 Vesicle exocytosis model

#### Input model

The vesicle exocytosis model describes coarse-grained spatiotemporal trajectories of vesicle exocytosis in pancreatic β-cells (dependent variables) after glucose stimulation, given an initial cell configuration (independent variables) (Fig. S2) (87).

**Fig. S3.**
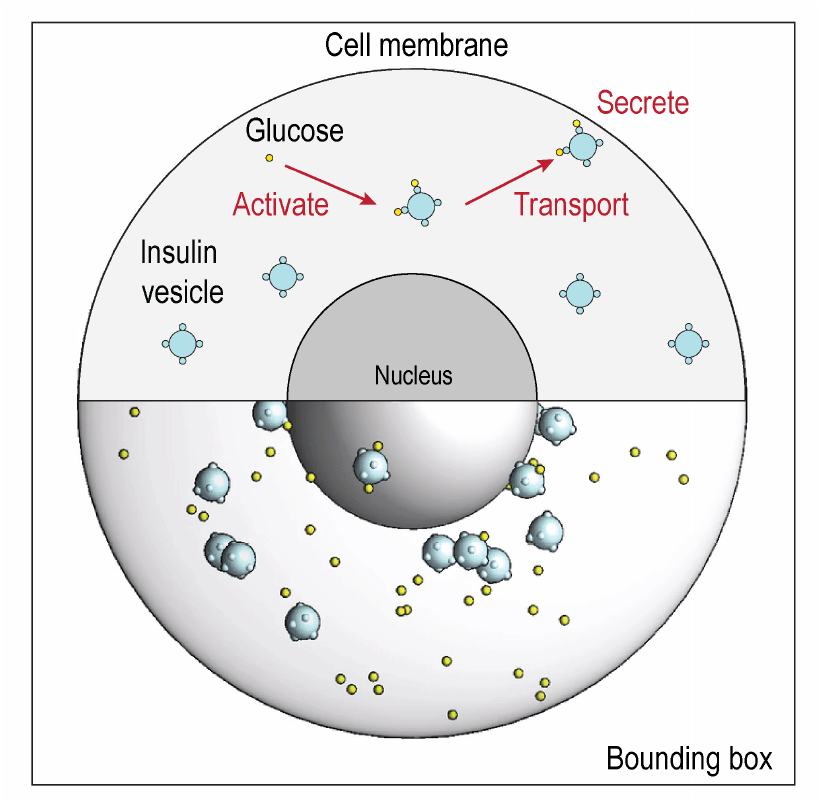
The vesicle exocytosis input model. The scheme indicates model components (top) and a snapshot from an actual coarse-grained Brownian dynamics simulation trajectory (bottom) (87). The model includes the cell membrane (light gray sphere), the nucleus (dark gray sphere), hundreds of insulin vesicles (light blue spheres), and thousands of glucose molecules (yellow spheres). The components are rescaled for visualization. Brownian dynamics simulations are restrained by various experimental data, including soft X-ray tomograms of the cell (19). 48 different 200 ms trajectories were computed for each of the following three treatment conditions (Table S3): (i) no glucose stimulation, (ii) 25 mM glucose stimulation, and (iii) 25 mM glucose stimulation with 10 nM Exendin-4 treatment. Exendin-4 is a peptide agonist of the glucagon-like peptide (GLP) receptor that attenuates postprandial plasma glucose (23).

**Table S3.**
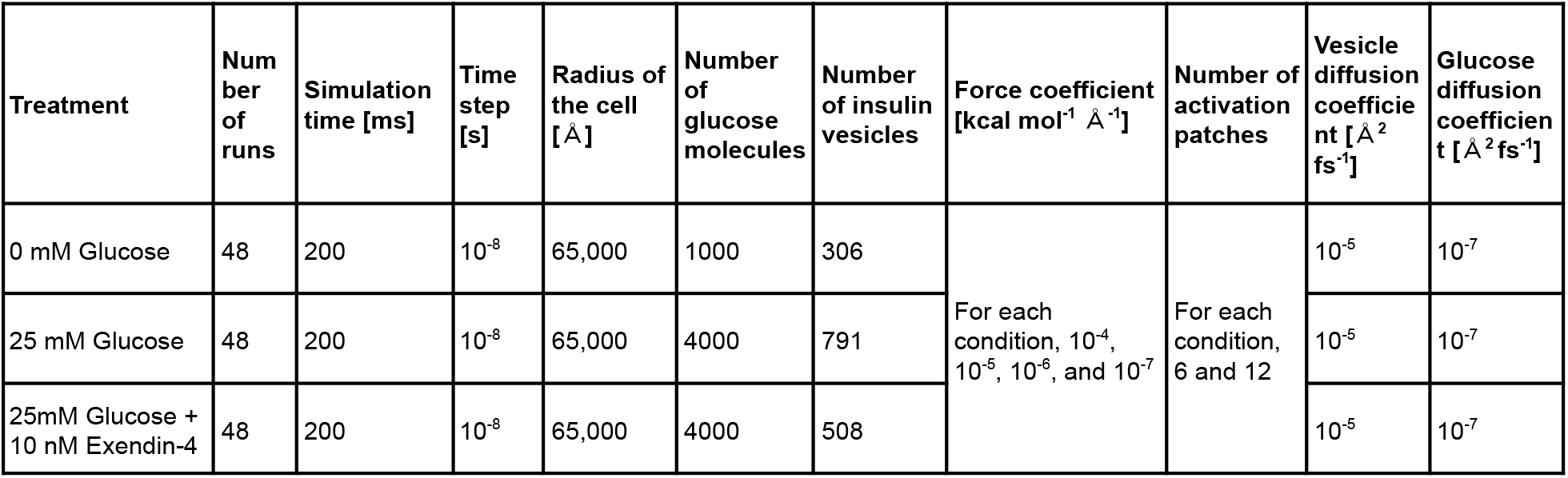
Model parameters of data-driven Brownian dynamics simulations under different conditions.

#### Surrogate model

The rationale behind the construction of the vesicle exocytosis surrogate model is described in Results (first example for Step 1 of metamodeling). This surrogate model includes a subset of the input model variables and new variables that are computed from input model variables. For example, total insulin secretion rate in the surrogate model is computed from insulin vesicle coordinates; these coordinates are in turn omitted from the surrogate model for practical reasons of dimensionality reduction. The conditional PDF parameters are fitted manually to the Brownian dynamics simulations to recapitulate the insulin secretion rates of the β-cell for different simulation conditions. The time step in the surrogate model (1 min) is longer than the entire time scale of the vesicle exocytosis model (200 ms); thus, a single time slice in the DBN of the surrogate model represents a single simulation trajectory of vesicle exocytosis. The surrogate vesicle exocytosis model simplifies the corresponding input model in three ways. First, the cell is secreting at a constant rate across one minute. Second, the surrogate model interpolates a linear Gaussian relationship from a discrete set of treatment conditions. Third, the surrogate model describes instantaneous rather than second phase insulin secretion occuring 30 minutes after glucose stimulation.

**Table S4.** Random variables, corresponding conditional PDFs, and conditional PDF parameters in the vesicle exocytosis surrogate model.

| Time-dependent variables and their conditional PDFs: |  |  |  |  |  |
| --- | --- | --- | --- | --- | --- |
| Name | Description | DBN time slice* [min] | Mean | Std-dev | Unit |
| $G^{VE}$ | Intracellular glucose concentration | $t_0$ | 2.55 | 0.1 | mM |
| | | $t + \Delta t$ | 2.55 | 0.01 | |
| $k_t^{VE}$ | Effective rate of vesicle trafficking towards the cellular periphery | $t_0$ | $5.0 + \alpha^{VE} S^{VE}(t_0)/2$ | 1.0 | $\text{m s}^{-1}$ |
| | | $t + \Delta t$ | $5.0 + \alpha^{VE} S^{VE}(t + \Delta t)/2$ | 0.1 | |
| $N_v^{VE}$ | Number of insulin vesicles in one β-cell | $t_0$ | $\beta^{VE} S^{VE}(t_0)$ | 1 | - |
| | | $t + \Delta t$ | $\beta^{VE} S^{VE}(t + \Delta t)$ | 0.1 | |
| $N_{patch}^{VE}$ | Number of activation patches per vesicle | $t_0$ | 6 | 0.1 | - |
| | | $t + \Delta t$ | $1.0 N_{patch}^{VE}(t)$ | 0.01 | |
| $N_{ins}^{VE}$ | Amount of insulin in one vesicle | $t_0$ | $1.8 \times 10^{-6}$ | $10^{-7}$ | pmol |
| | | $t + \Delta t$ | $1.0 N_{ins}^{VE}(t)$ | $10^{-8}$ | |
| $S^{VE}$ | Insulin secretion rate of one β-cell | $t_0$ | $9.32 \times 10^{-9}$ | $10^{-10}$ | $\text{pM min}^{-1}$ |
| | | $t + \Delta t$ | $k_G^{VE} G^{VE}(t) + k_p^{VE} N_{patch}^{VE}(t + \Delta t) + k_{ins}^{VE} N_{ins}^{VE}(t + \Delta t) + k_D^{VE} D_v^{VE}(t + \Delta t) + k_R^{VE} R_{cell}^{VE}(t + \Delta t)$ | $10^{-11}$ | |
| Time-independent variables represented as conditional PDF parameters: |  |  |  |  |  |
| $R_{cell}^{VE}$ | Radius of the β-cell | t | 6 | 0.1 | $\mu\text{m}$ |
| $D_v^{VE}$ | Diffusion coefficient of insulin vesicles in the beta cell | t | 0.0032 | $10^{-4}$ | $\text{\AA}^2 \text{fs}^{-1}$ |

| Parameters of conditional PDFs: |  |  |  |  |  |
| --- | --- | --- | --- | --- | --- |
| Name | Description | DBN time slice* [min] | Mean | Std-dev | Unit |
| $\alpha^{VE}$ | Correlation between insulin secretion rate and the force coefficient of vesicle transport | - | $1.073 \times 10^9$ | - | $\text{m s}^{-1} \text{ pM}^{-1} \text{ min}$ |
| $\beta^{VE}$ | Correlation between insulin secretion rate and the number of insulin vesicles in the cell | - | $3.219 \times 10^{10}$ | - | $\text{pM}^{-1} \text{ min}$ |
| $k_p^{VE}$ | Coefficient for the number of activation patches on the vesicle surface accelerating insulin secretion rate | - | $2.55 \times 10^{-10}$ | - | $\text{pM min}^{-1}$ |
| $k_G^{VE}$ | Coefficient for the intracellular glucose concentration stimulating the insulin secretion | - | $3.4 \times 10^{-9}$ | - | $\text{pM min}^{-1} \text{ mM}^{-1}$ |
| $k_{ins}^{VE}$ | Coefficient for the number of insulin molecules in each vesicle determining insulin secretion rate | - | 0.01 | - | $\text{pM min}^{-1}$ |
| $k_D^{VE}$ | Coefficient for the vesicle diffusion promoting insulin secretion rate | - | $10^{-7}$ | - | $\text{pM min}^{-1} \text{ A}^{-2} \text{ fs}$ |
| $k_R^{VE}$ | Coefficient for the cell radius reducing insulin secretion rate | - | $-3.2 \times 10^{-9}$ | - | $\text{pM min}^{-1} \mu\text{m}^{-1}$ |

\* $t_0$ is the initial time slice in the DBN; $t + \Delta t$ is the time slice that follows time slice $t$ by a time step of $\Delta t$ .

### 1.4 GSIS signaling model

#### Input model

The GSIS signaling model describes the dynamics of the molecular signaling network leading to insulin secretion in β-cells following glucose stimulation and a GLP1 hormone signal (Fig. S4). The model computes insulin secretion rates as well as concentrations of various signaling molecules, such as ATP, cAMP, and Ca^2+^ over time (dependent variables), for a given starting condition (independent variables), using a system of linear ODEs.

**Figure S4.**
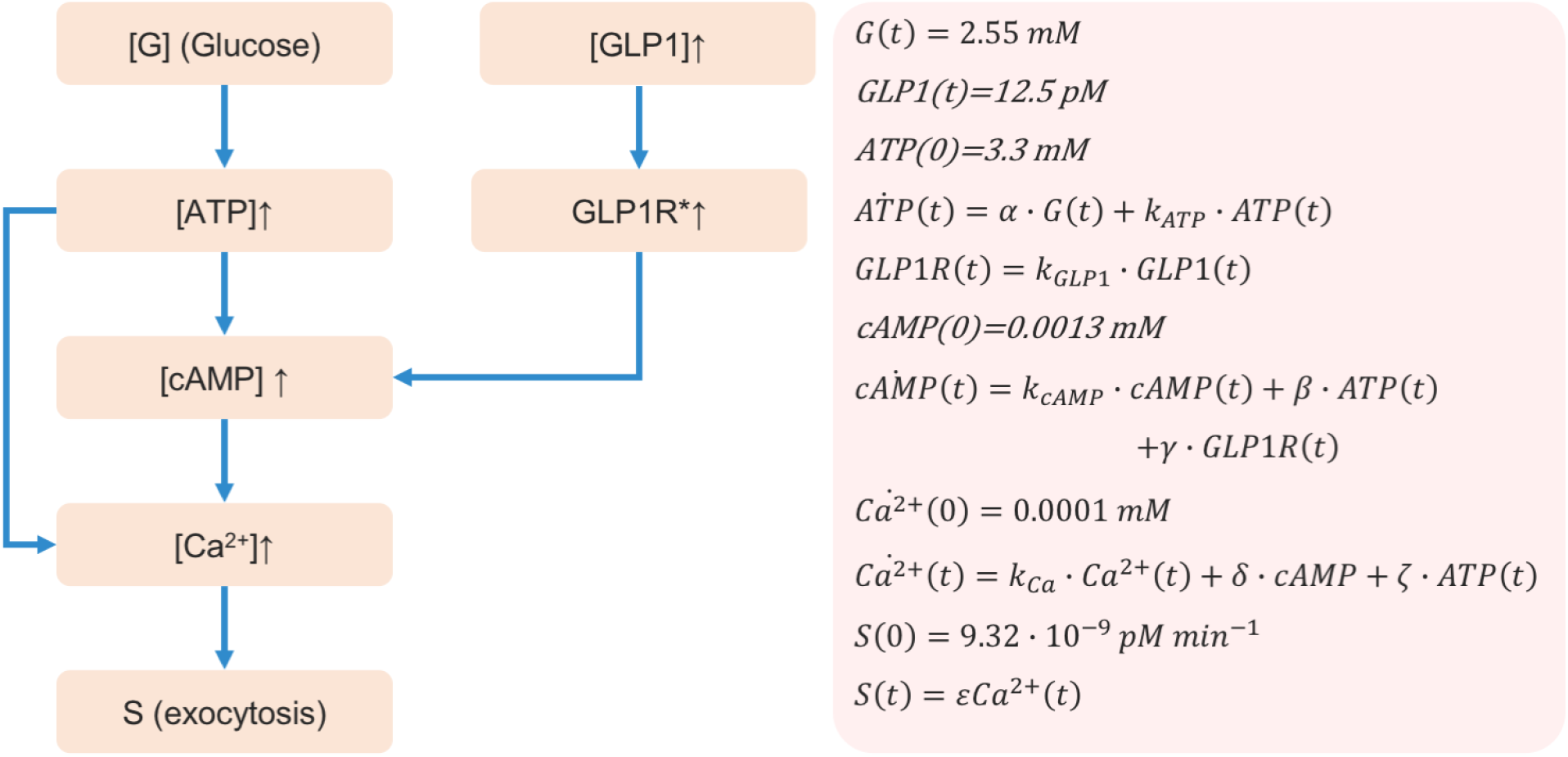
The GSIS signaling input model. The signaling network topology (left) is a combination of the linear pathways leading from glucose stimulation to insulin secretion, based on pathway hsa04911 in the KEGG database (88) and the signaling pathway leading to cAMP-dependent enhancement of insulin secretion following activation of the GLP1R receptor by the peptide hormone GLP1. GLP1R* is a variable that indicates GLP1R relative activity levels. Linear ODEs (right) describe the time evolution of the network component, leading to insulin secretion, *S(t)*. ODE coefficient values are identical to the values of the corresponding parameters of the conditional PDFs (Table S5). The following simplifying assumptions were made during model construction: (1) the system dynamics are described by linear equations; (2) the signaling network of pancreatic β cells is simplified, for example, by merging alternative pathways through which cAMP and ATP modulate calcium release from various cellular and extracellular compartments and omitting feedback loops in the network; and (3) identical parameters are used for both healthy and T2D subjects.

#### Surrogate model

The DBN describing the GSIS signaling surrogate model is a discretized, probabilistic version of the linear ODEs of the corresponding input model (Fig. 2, Table S5). As with other surrogate models, standard deviations were added to reflect our uncertainty in variable values.

**Table S5.**
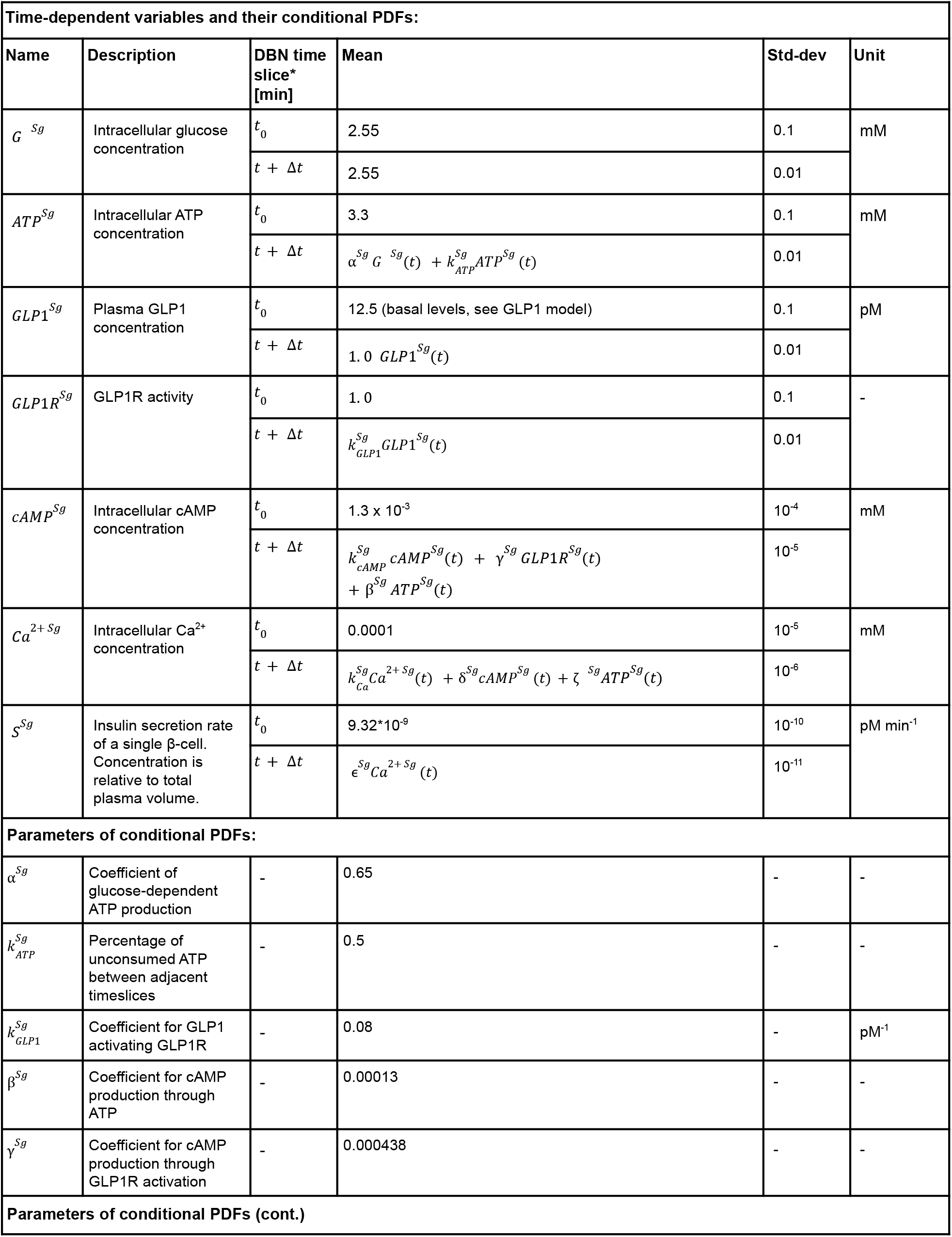
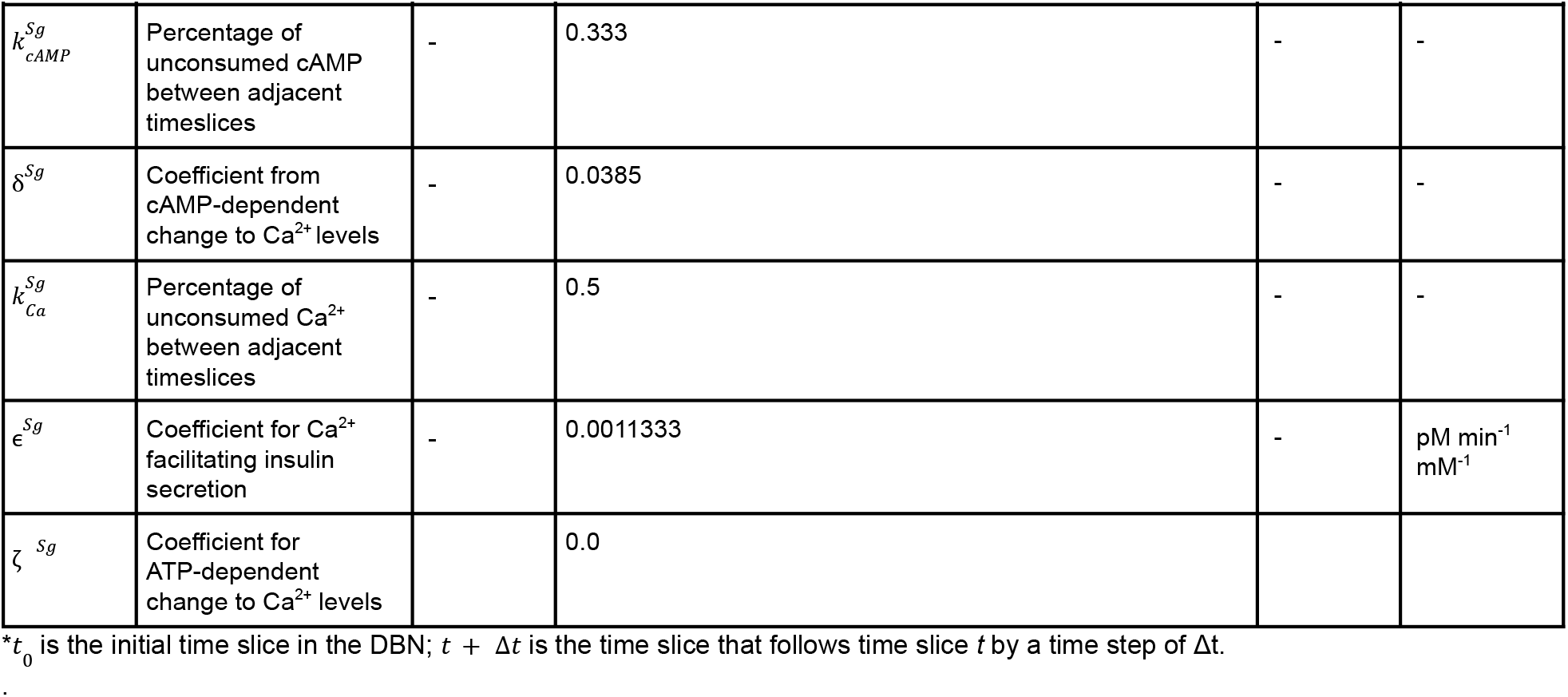
Random variables, corresponding conditional PDFs, and conditional PDF parameters in the GSIS signaling surrogate model.

### 1.5 Insulin metabolism model

#### Input model

The insulin metabolism model predicts activation of cellular metabolic pathways (dependent variables) for different treatment conditions (independent variables), based on experimental measurements of metabolomic signatures obtained using liquid chromatography-mass spectrometry.

**Figure S5.**
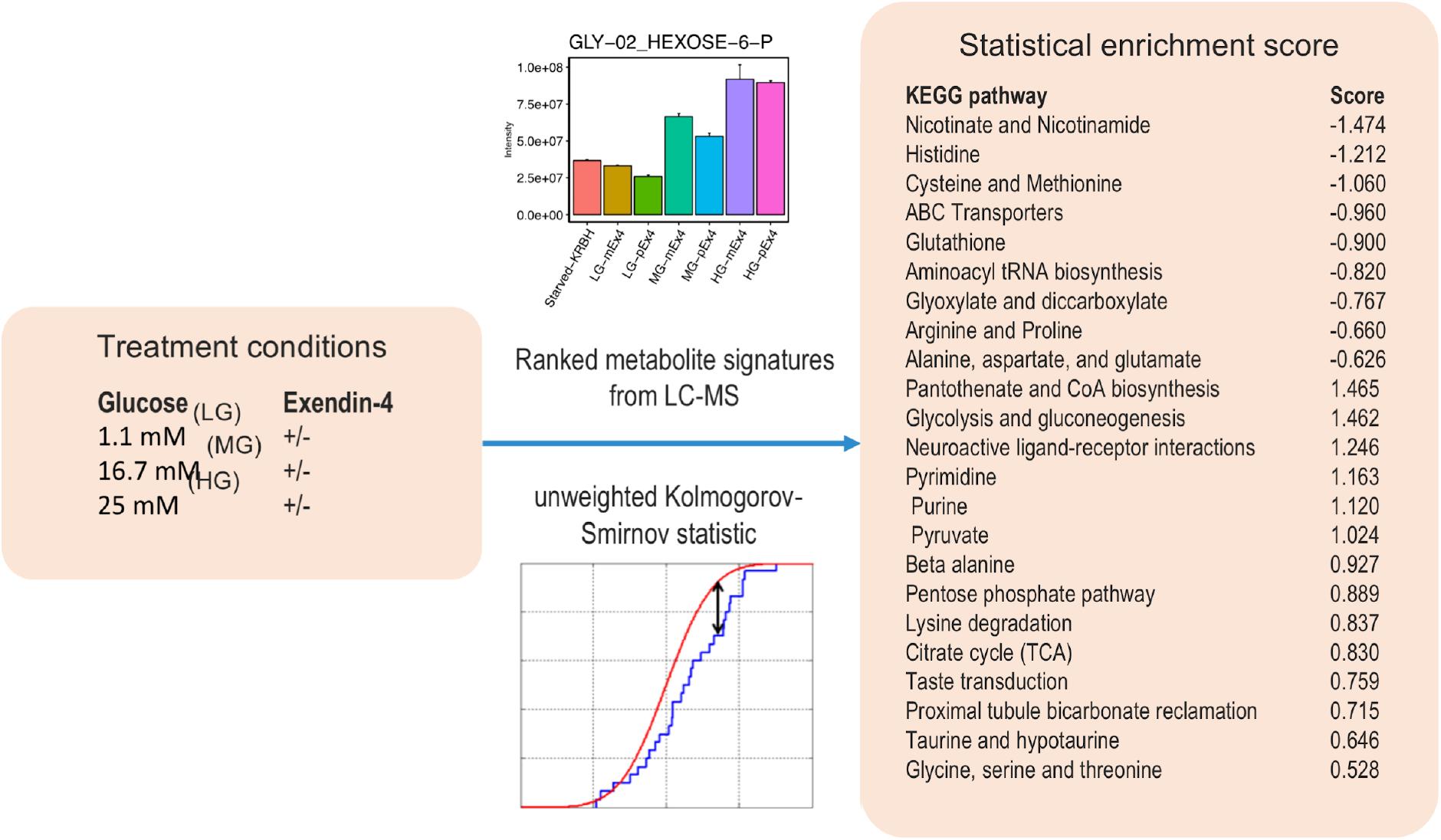
The insulin metabolism input model. Enrichment analysis for functional metabolic pathways is informed by metabolite quantities measured by liquid chromatography-mass spectrometry (LC-MS) measurements on INS-1E cells under different glucose and Exendin-4 treatments. Based on these measurements, a statistical model was constructed that outputs normalized enrichment scores for 25 different metabolic pathways from the KEGG pathways database (116), in response to different treatment conditions. Metabolomic signatures were ranked based on log2-transformed fold change observed for a given perturbation (*e*.*g*., LG/MG/HG co-stimulated with Exendin-4 compared with incretin-free LG/MG/HG treated INS-1E cells). Metabolites were annotated using KEGG COMPOUND ID (*e*.*g*., D-Glucose: C00031). The enrichment analyses were run with unweighted Kolmogorov-Smirnov statistic using the Broad Institute’s GSEA java applet against a library containing all KEGG metabolic pathways. Normalized enrichment scores (NES) were calculated to determine if metabolic pathways were overrepresented at the top or the bottom of the given rank lists. Statistical significance scores were assessed by 5,000 permutations of the ranked lists. The permutation-based P-value is defined by the fraction of randomly permuted lists resulting in the NES values greater than or equal to the observed NES. The KEGG metabolic pathway library (c2_kegg_gene_cpd_set.gmt) was constructed by scraping the KEGG API. NES scores for each KEGG pathway at each treatment conditions are provided in the Excel file https://github.com/salilab/metamodeling/blob/master/data/072919-INS1e-30min-Enrichment-analysis-cleaned-summary.xlsx.

#### Surrogate model

The insulin metabolism surrogate model describes a parametrized linear relation between the treatment conditions (the independent variables), the TCA cycle enrichment score (one of the dependent variables), and the concentrations of intracellular ATP (an additional variable estimated from enrichment of TCA). The insulin metabolism surrogate model simplifies the corresponding input model in three ways. First, it summarizes a narrow aspect of the input model that is of particular interest. Second, it interpolates a linear Gaussian relationship from a discrete set of treatment conditions. Third, it assumes that the change in TCA cycle activation and ATP concentration occurs instantaneously rather than 30 minutes after glucose stimulation. The model parameters were manually fitted to reproduce the empirical relation between independent and dependent variables in the corresponding input model. While this model describes INS-1e cells rather than primary β-cells, differences in pathway enrichment among cell types are accounted for during the coupling stage (Discussion).

**Table S6.**
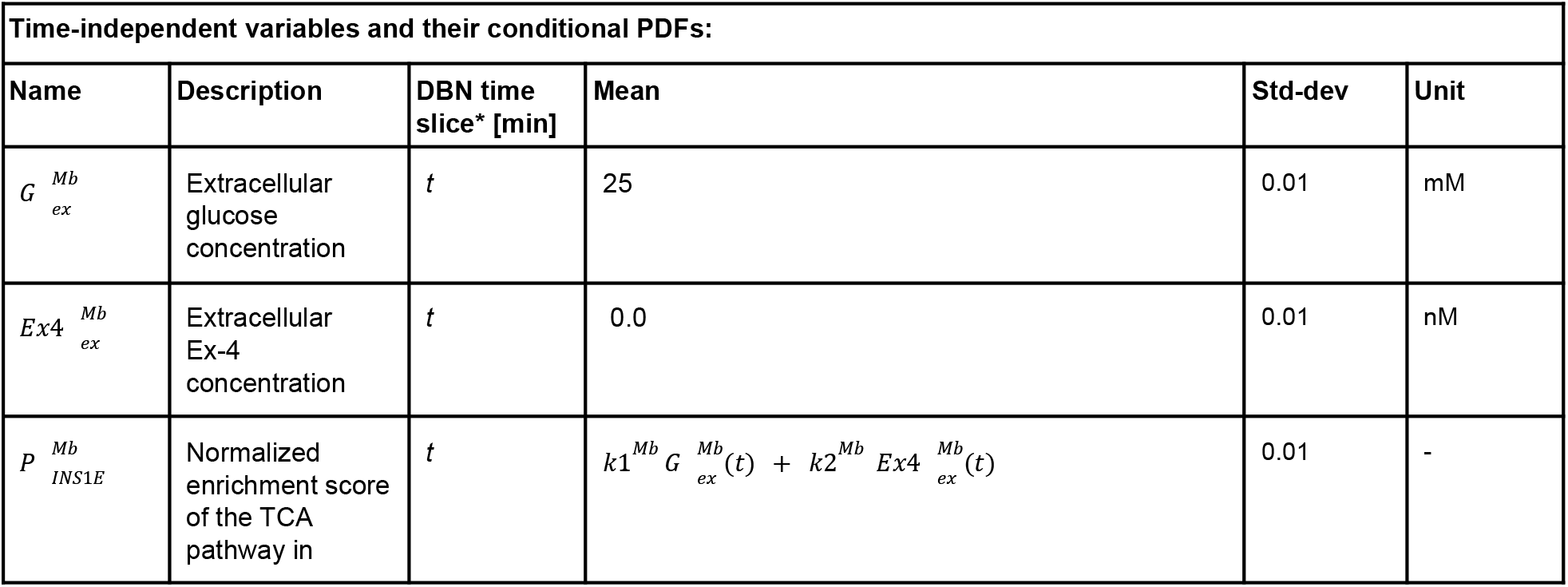
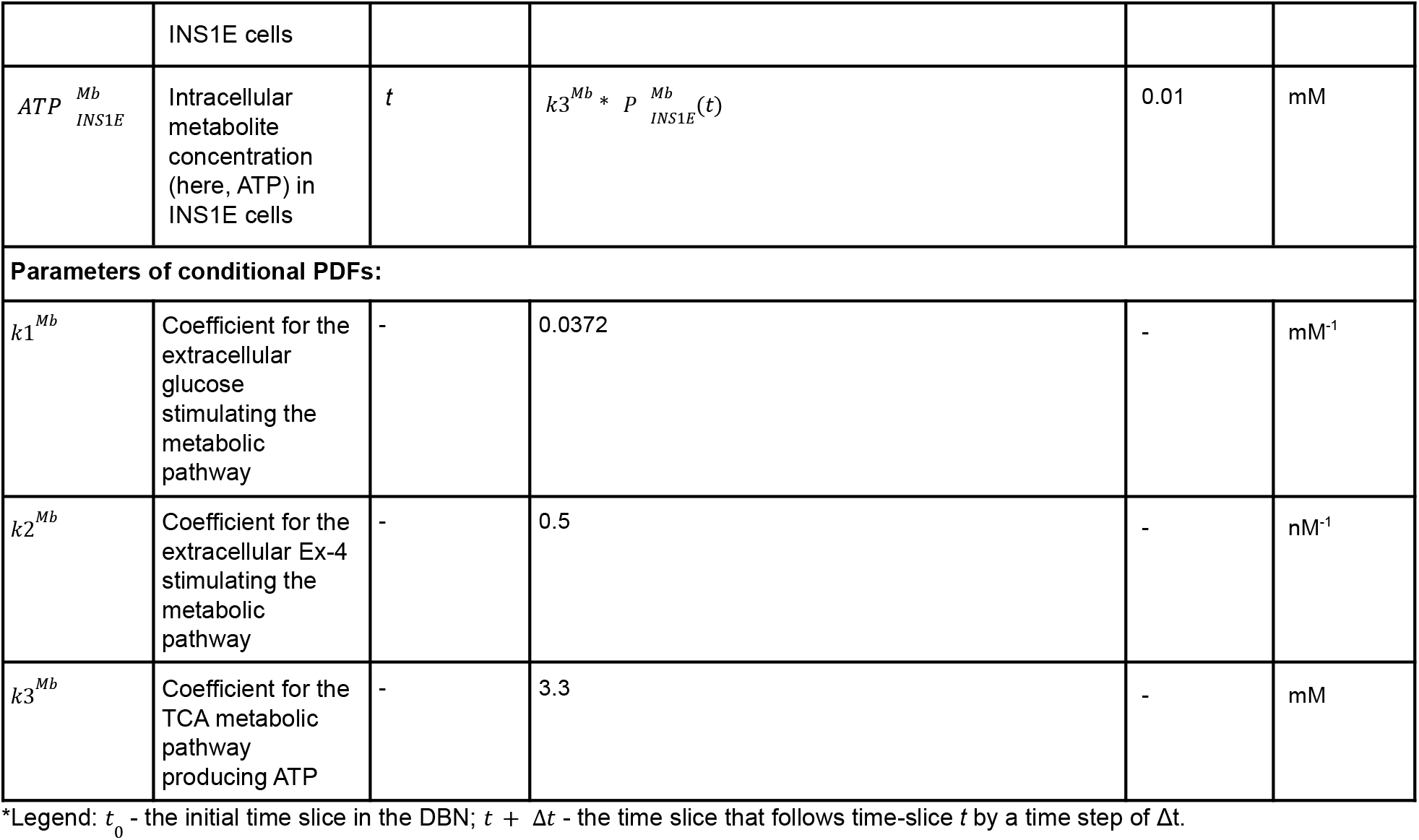
Random variables, corresponding conditional PDFs, and conditional PDF parameters in the insulin metabolism surrogate model.

#### Cell cultures

INS-1E *Rattus Norvegicus* insulinoma cells were obtained from the Cell Culture Core of the Raymond Stevens lab (Bridge Institute USC). Cells were maintained in modified RPMI 1640 supplemented with 5% horse serum, 1 mM sodium pyruvate, 50 μM β-mercaptoethanol, 2 mM glutamine, and 10 mM HEPES in monolayer. All cells were grown in a humidified incubator at 5% CO_2_ and 37°C. They were used between 30-50 passages of thawing. Cell counting and viability were assessed using trypan blue staining with a TC20 automated cell counter (BioRad).

#### Cell stimulation and intracellular metabolites extraction

Cells were plated on 6-well plates at a density of 7,000 cells/cm^2^. When cell densities reached 70%, the media was removed, cells were washed twice with 2 mL of PBS, followed by adding 5 mL of KRBH buffer with 0 mM glucose to cells. The cells were starved for 30 min prior to treatment. Following starvation, KRBH buffers were removed and the cells were treated with 1.1 mM, 16.7 mM, and 25 mM of glucose without or with 10 nM of Exendin-4. After 30 min, supernatants were collected, spun-down, and assayed using Mercodia Rat Insulin ELISA kit (10-1250-01) according to the manufacturer protocol. The culture plates were cooled on ice, and the cells were washed with 1 mL of cold ammonium acetate. The methanol cell suspensions were scraped and transferred to Eppendorf tubes, followed by centrifugation at 4°C. The supernatants were transferred to new Eppendorf tubes, and the pellets were re-extracted with another 350 μL of -80°C methanol. The second methanol extraction was spun-down, and the supernatants were pooled with the first extraction. Metabolites were speed-vac’ed to dryness, resuspended in LC-MS grade water, and submitted to LC-MS.

#### Liquid chromatography-mass spectrometry (LC–MS) metabolomics

Samples were randomized and analyzed on a Q Exactive Plus hybrid quadrupole-Orbitrap mass spectrometer coupled to an UltiMate 3000 UHPLC system (Thermo Scientific). The mass spectrometer was run in polarity switching mode (+3.00 kV/-2.25 kV) with an m/z window ranging from 65 to 975. Mobile phase A was 5 mM NH4AcO, *p*H 9.9, and mobile phase B was acetonitrile. Metabolites were separated on a Luna 3 µm NH2 100 Å (150 × 2.0 mm) column (Phenomenex). The flow rate was 300 µL/min, and the gradient was from 15% A to 95% A in 18 min, followed by an isocratic step for 9 min and re-equilibration for 7 min. All samples were run in biological triplicate. Metabolites were detected and quantified by the area-under-the-curve, based on retention time and accurate mass (≤ 8 ppm) using the TraceFinder 3.3 (Thermo Scientific) software. Intracellular data was normalized to the cell number at the time of extraction.

### 1.6 Virtual screening model

#### Input model

The virtual screening model describes the increase in the activity level of GLP1R (dependent variable) for different concentrations of various GLP1 agonist compounds, based on their rank in a library of compounds (independent variables).

**Figure S6.**
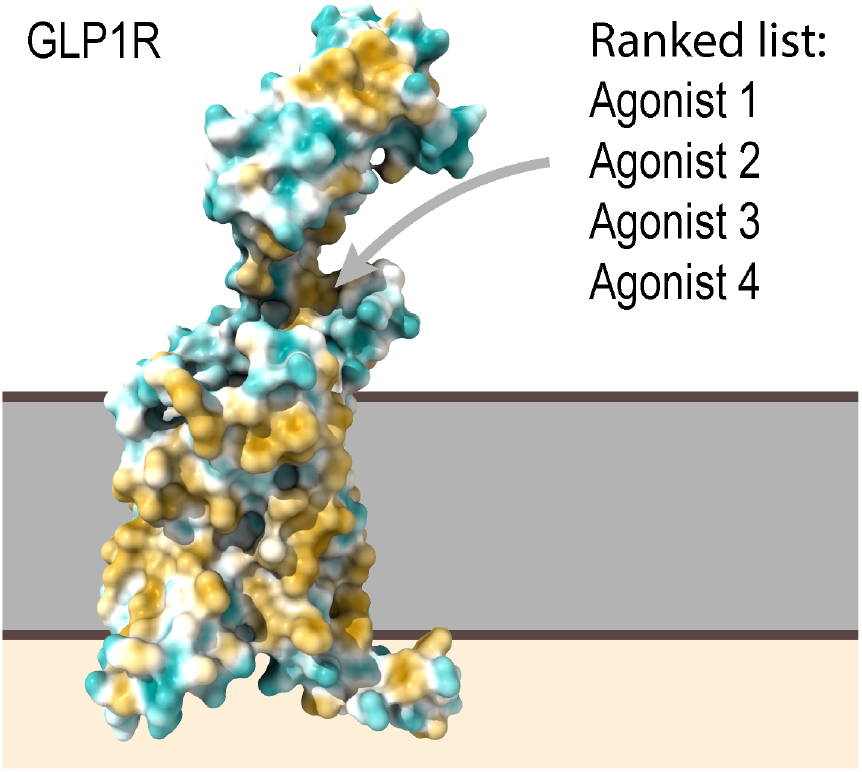
The virtual screening input model. The model is based on computational docking of 5,689 potential agonists against an atomic structure of GLP1R (93). These potential agonists were ranked by their predicted affinity to the allosteric site of GLP1R. We selected the top four compounds. The model computes the increase in GLP1R activity as a function of the rank of the compound *A* and its concentration 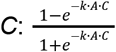, where maximal relative activation *F* is set to 310.0 fold and a concentration normalization coefficient *k* was set to 1.1 pM^-1^.

#### Surrogate model

The virtual screening surrogate model is a linear approximation of the corresponding input model, obtained through manual fitting. It simplifies the corresponding input model in two ways: First, activity is related to the rank and concentration *via* a linear Gaussian. Second, identical variable and parameter values are used for both healthy and T2D subjects.

**Table S7.**
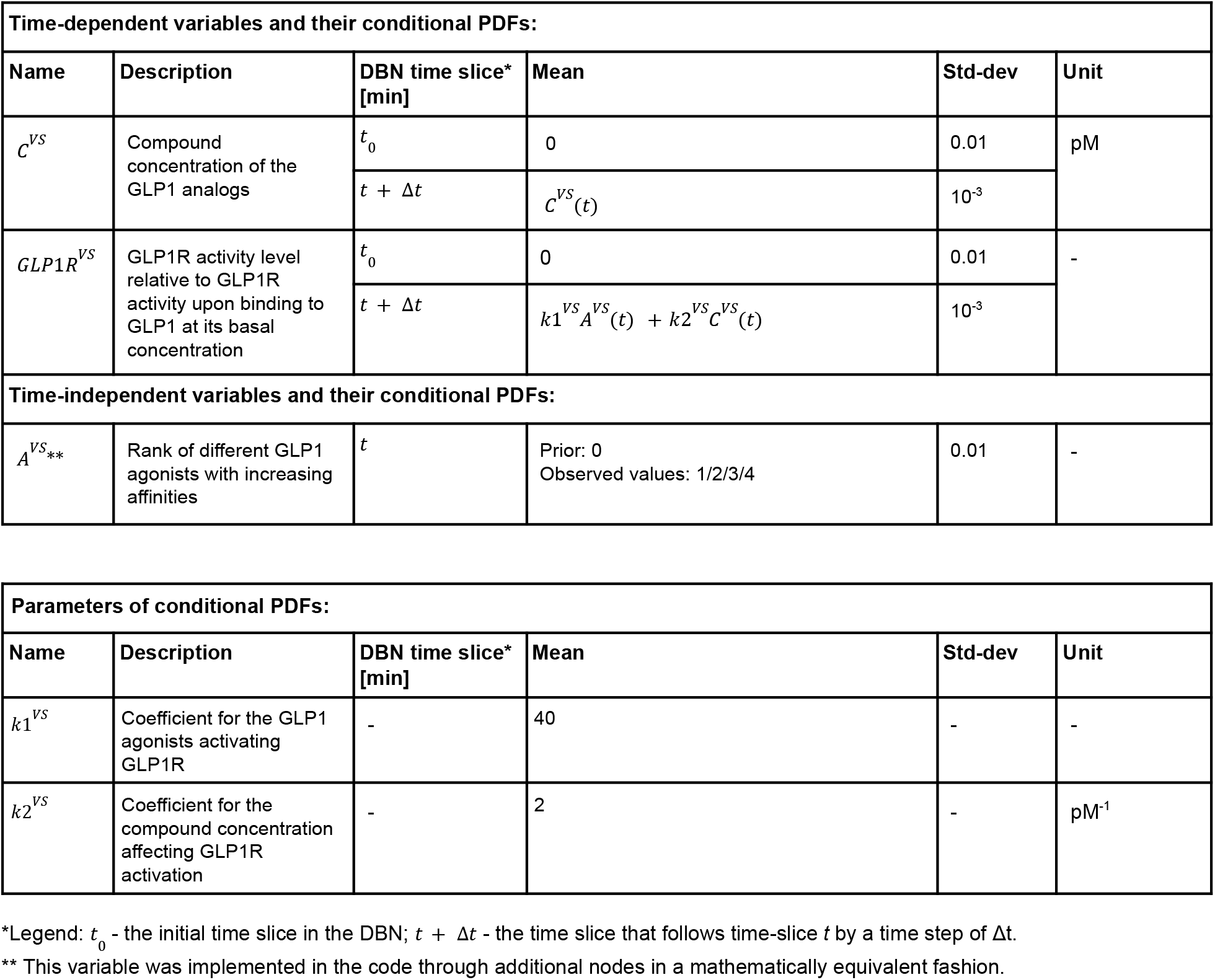
Random variables, corresponding conditional PDFs, and conditional PDF parameters in the virtual screening surrogate model.

### 1.7 Glucose intake data model

*Input model*. The GI data model tabulates data on the rate of glucose intake from food digestion, 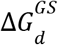 (Table S8). The GI data model illustrates how real-world data can be integrated through metamodeling and coupled with other models.

**Table S8.** GI data model. The data was simulated at 1 min interval using a sigmoid postprandial response model. The postprandial response model itself is in turn based on empirical data that are typically measured by glucose sensors as the rate of appearance from the intestine (20).

| Time<br>[min] | $\Delta G_d$<br>[mM<br>min<sup>-1</sup>] |
| --- | --- |
| 0 | 0.047 |
| 1 | 0.057 |
| 2 | 0.069 |
| 3 | 0.083 |
| 4 | 0.099 |
| 5 | 0.118 |
| 6 | 0.140 |
| 7 | 0.166 |
| 8 | 0.195 |
| 9 | 0.226 |
| 10 | 0.261 |
| 11 | 0.298 |
| 12 | 0.337 |
| 13 | 0.375 |
| 14 | 0.411 |
| 15 | 0.443 |
| 16 | 0.470 |
| 17 | 0.489 |
| 18 | 0.498 |
| 19 | 0.498 |
| 20 | 0.489 |
| 21 | 0.470 |
| 22 | 0.443 |
| 23 | 0.411 |
| 24 | 0.375 |
| 25 | 0.337 |
| 26 | 0.298 |
| 27 | 0.261 |
| 28 | 0.226 |
| 29 | 0.195 |
| 30 | 0.166 |
| 31 | 0.140 |
| 32 | 0.118 |
| 33 | 0.099 |
| 34 | 0.083 |
| 35 | 0.069 |
| 36 | 0.057 |
| 37 | 0.047 |
| 38 | 0.039 |
| 39 | 0.032 |
| 40 | 0.026 |
| 41 | 0.022 |
| 42 | 0.018 |
| 43 | 0.015 |
| 44 | 0.012 |
| 45 | 0.010 |
| 46 | 0.008 |
| 47 | 0.007 |
| 48 | 0.005 |
| 49 | 0.004 |
| 50 | 0.004 |
| 51 | 0.003 |
| 52 | 0.002 |
| 53 | 0.002 |
| 54 | 0.002 |
| 55 | 0.001 |
| 56 | 0.001 |
| 57 | 0.001 |
| 58 | 0.001 |
| 59 | 0.000 |
| 60 | 0.000 |

#### Surrogate model

The GI data surrogate model relies on the rate of glucose intake in the input model, with standard deviations reflecting data uncertainties.

**Table S9.** Conditional probability distributions for Gaussian variables in the GI data model.

| Variable name | Description | Values | Unit |
| --- | --- | --- | --- |
| $\Delta G_d^{GS}$ | Rate of glucose intake from food digestion | Table S8 | mM min<sup>-1</sup> |

### 1.8 GLP1 data model

#### Input model

The GLP1 data model defines classes for four discrete values of the plasma concentration of GLP1 (Table S10); the classification is the same for both normal and T2D patients.

**Table S10.**
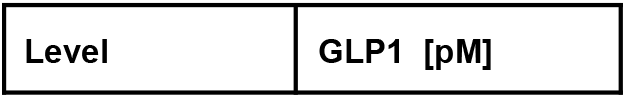
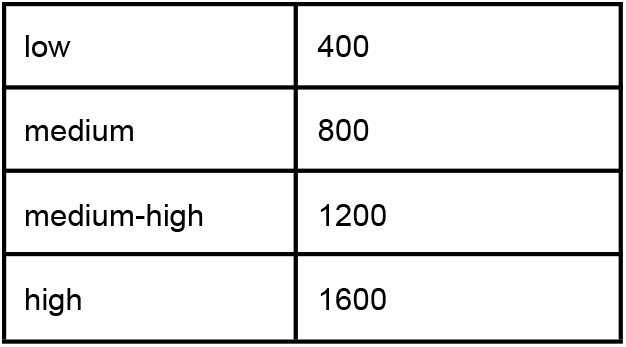
The GLP1 data model.

| Level | GLP1 [pM] |
| --- | --- |
| low | 400 |
| medium | 800 |
| medium-high | 1200 |
| high | 1600 |

#### Surrogate model

The GLP1 data surrogate model is the observation made in the corresponding model. It becomes probabilistic only during the coupling stage, in which a conditional PDF of the *GLP1*^*GL*^ variable relates it to a coupling variable *GLP1*^*C*^, as described in SI Appendix: Supplementary Text 2.

**Table S11.**
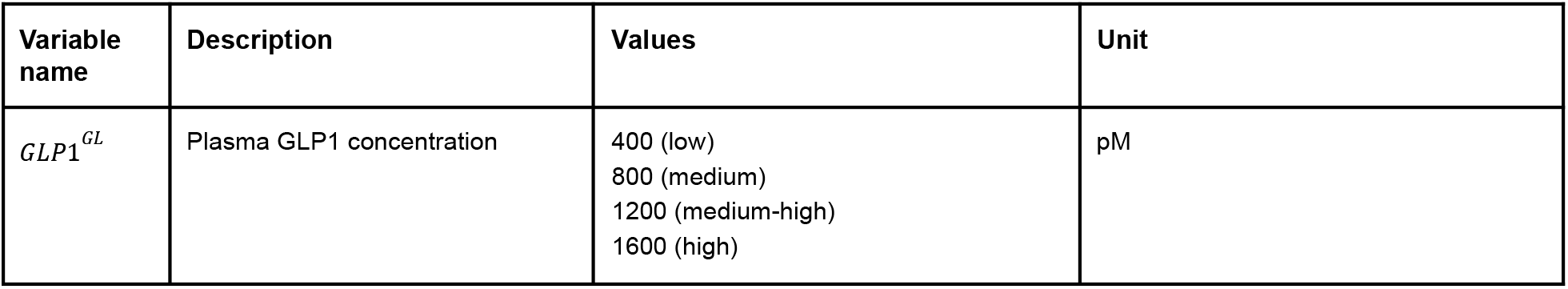
Conditional probability distributions for Gaussian variables in the GL data model.

| Variable name | Description | Values | Unit |
| --- | --- | --- | --- |
| $GLP1^{GL}$ | Plasma GLP1 concentration | 400 (low)<br>800 (medium)<br>1200 (medium-high)<br>1600 (high) | pM |

### 2. Coupling and coupling variables

In the coupling stage, coupling variables were introduced as random variables with linear Gaussian conditional PDFs, relating them to variables from surrogate models (Table S10). In addition, the conditional PDFs of some variables from surrogate models were modified to include dependencies on coupling variables (Table S11). When variables from several models inform a coupling variable, our prior assigns even confidence to information from all variables. For instance, the coupling variable 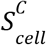 is statistically coupled to *S*^*VE*^ and *S*^*Sg*^ with equal weights. When additional prior information is available, it is used to inform the coupling. For example, intracellular glucose concentration has been estimated to be half of the plasma glucose concentration (95).

**Table S12.** Conditional PDFs for Gaussian variables of the coupling variables.

| Coupler | Description | Time slice, [min] | Mean | Std-dev | Unit |
| --- | --- | --- | --- | --- | --- |
| $G_{pl}^C$ | Plasma glucose concentration | t | $G^{PR}(t)$ | $10^{-3}$ | mM |
| $G_{cell}^C$ | Intracellular Glucose concentration | t | $G_{pl}^C(t)/2$ | $10^{-3}$ | mM |
| $GLP1R^C$ | GLP1R activity | t | $1.0 GLP1R^{VS}(t)$ | $10^{-4}$ | - |
| $ATP_{INS1E}^C$ | Intracellular ATP concentration | t | $1.0 \, ATP_{INS1E}^{Mb}(t)$ | $10^{-3}$ | mM |
| $S_{pa}^C$ | Insulin secretion rate of the pancrea | t | $S_{pa}^{Pa}(t)$ | $10^{-3}$ | pM min<sup>-1</sup> |
| $S_{cell}^C$ | Insulin secretion rate of one β-cell | t | $S^{Sg}(t)/2 + S^{VE}(t)/2$ | $10^{-11}$ | pM min<sup>-1</sup> |
| $\Delta G_d^C$ | Rate of glucose intake from food digestion | t | $1.0 \, \Delta G_d^{PR}$ | $10^{-6}$ | mM min<sup>-1</sup> |
| $GLP1^C$ | Plasma GLP1 concentration | t | $1.0 \, GLP1^{Sg}(t)$ | $10^{-3}$ | pM |

**Table S13.** Modified conditional PDFs for variables in surrogate models that become dependent on coupling variables after the coupling stage.

| Variable name | Description | Time slice, [min] | Mean | Std-dev | Unit |
| --- | --- | --- | --- | --- | --- |
| $S^{PR}$ | Pancreatic insulin secretion rate | $t + \Delta t$ | $(Y^{PR}(t + \Delta t) + K^{PR} \Delta G_d^{PR}(t) + 1.0 S_b^{PR}(t + \Delta t))/3 + 2/3 S_{pa}^C(t + \Delta t)$ | $10^{-3}$ | pM min<sup>-1</sup> |
| $S_{cell}^{Pa}$ | Insulin secretion of a cell | $t$ | $S_{cell}^C(t)$ | $10^{-10}$ | pM min<sup>-1</sup> |
| $G^{VE}$ | Basal intracellular glucose concentration | $t$ | $1.0 G_{cell}^C(t)$ | $10^{-3}$ | mM |
| $G^{Sg}$ | Intracellular glucose concentration | $t$ | $1.0 G_{cell}^C(t)$ | $10^{-3}$ | mM |
| $ATP^{Sg}$ | Intracellular ATP concentration | $t + \Delta t$ | $0.9 (\alpha^{Sg} G_{ic}^{Sg}(t) + k_{ATP}^{Sg} ATP^{Sg}(t)) + 0.11 ATP_{INS1E}^C(t + \Delta t)$ | $10^{-4}$ | mM |
| $GLP1R^{Sg}$ | GLP1R activity | $t$ | $k_{GLP1}^{Sg} GLP1^{Sg}(t) + GLP1R^C(t)$ | $10^{-3}$ | - |
| $\Delta G_d^{GI}$ | Rate of glucose intake from food digestion | $t$ | $1.0 \Delta G_d^C(t)$ | $10^{-6}$ | mM min<sup>-1</sup> |
| $GLP1^{GL}$ | Plasma GLP1 concentration | $t$ | $1.0 GLP1^C(t)$ | $10^{-3}$ | pM |

## Additional Supplementary Figures and Movie

**Figure S14.**
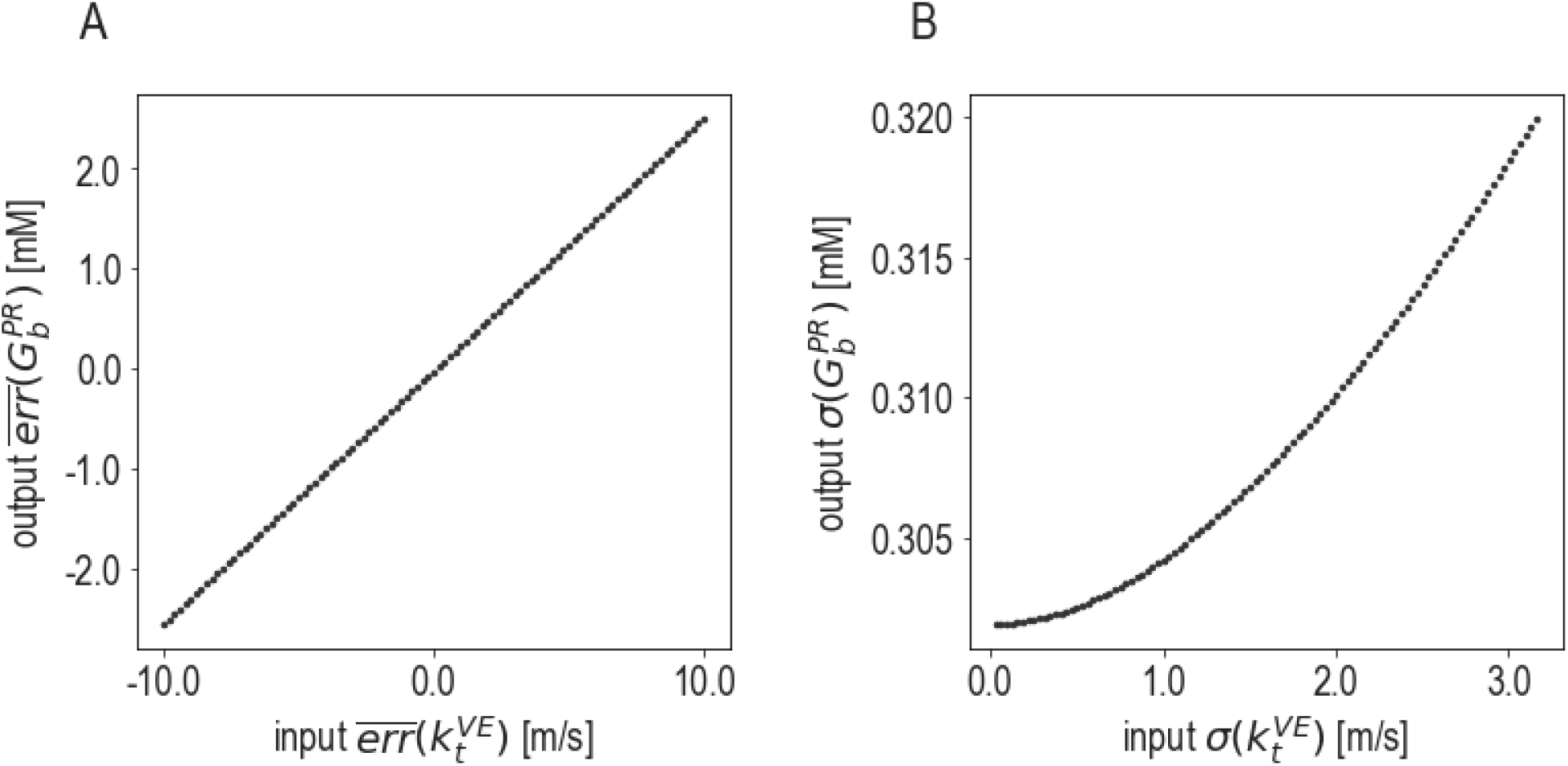
Effect of metamodeling on model accuracy and precision. (A) Statistical dependency of the output systematic error 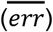 of the variable 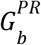 in the postprandial response model on the input systematic error of the variable 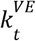 in the vesicle exocytosis model. The input systematic error for 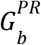 is 0.0. The coupling coefficient corresponds to the slope of the line. (B) Statistical dependency of the output random error (*σ*) of 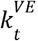 on the input random error of 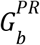. Input 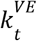 was presented as evidence with equal values for all time steps. The input random error for 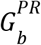 is 1.0. For both (A) and (B), the reference values used for computing the systematic and random errors of 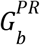 and 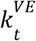 are 5.1 mM and 10.0 m/s, respectively. All output values are at *t* = 100 min.

**Movie S1.**
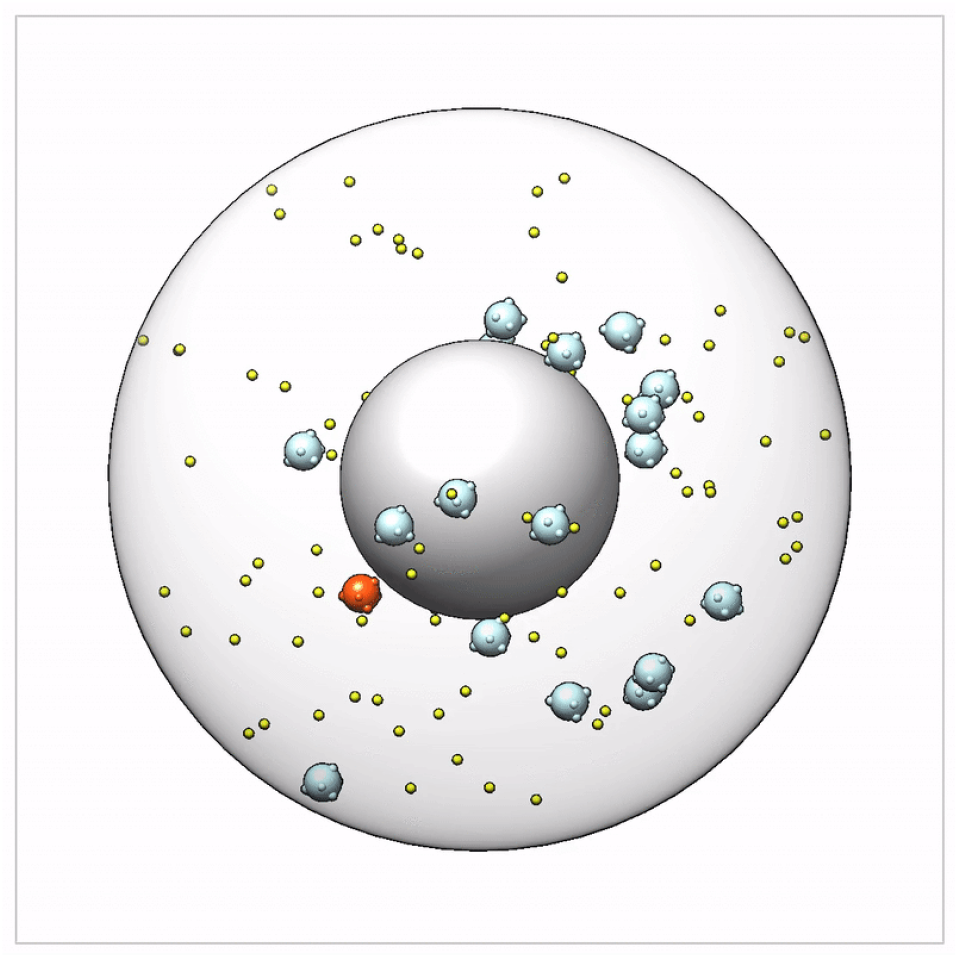
A sample coarse-grained Brownian dynamics trajectory contributing to the vesicle exocytosis input model. A 32.8 μs trajectory is shown. See legend of Figure S3. A single insulin vesicle is colored in red, highlighting the spatiotemporal trajectory of a single secretion event, including glucose activation, transport, and secretion. Following secretion, the vesicle is “reborn” inside the cytoplasm, modeling a new vesicle biogenesis event. The simulation time step is 10 nanoseconds, and the animation shows snapshots at 200 nanosecond intervals.

## Notes

http://github.com/salilab/metamodeling

### Competing Interest Statement

The authors have declared no competing interest.

## References

1. David Sadave, David M Hillis, H Craig Heller, May R Berenbaum, Ed., Life the Science of Biology (W. H. Freeman; 10th edition, 2014).

2. A. Warshel, M. Karplus, Calculation of ground and excited state potential surfaces of conjugated molecules. I. Formulation and parametrization. J. Am. Chem. Soc. 94, 5612–5625 (1972).

3. A. Warshel, M. Levitt, Theoretical studies of enzymic reactions: dielectric, electrostatic and steric stabilization of the carbonium ion in the reaction of lysozyme. J. Mol. Biol. 103, 227–249 (1976).

4. M. P. Rout, A. Sali, Principles for Integrative Structural Biology Studies. Cell 177, 1384–1403 (2019).

5. S. J. Kim, J. Fernandez-Martinez, I. Nudelman, Y. Shi, W. Zhang, B. Raveh, T. Herricks, B. D. Slaughter, J. A. Hogan, P. Upla, I. E. Chemmama, R. Pellarin, I. Echeverria, M. Shivaraju, A. S. Chaudhury, J. Wang, R. Williams, J. R. Unruh, C. H. Greenberg, et al., Integrative structure and functional anatomy of a nuclear pore complex. Nature 555, 475–482 (2018).

6. S. R. McGuffee, A. H. Elcock, Diffusion, crowding & protein stability in a dynamic molecular model of the bacterial cytoplasm. PLoS Comput. Biol. 6, e1000694 (2010).

7. M. Roy, S. D. Finley, Computational Model Predicts the Effects of Targeting Cellular Metabolism in Pancreatic Cancer. Front. Physiol. 8, 217 (2017).

8. Z. A. King, J. Lu, A. Dräger, P. Miller, S. Federowicz, J. A. Lerman, A. Ebrahim, B. O. Palsson, N. E. Lewis, BiGG Models: A platform for integrating, standardizing and sharing genome-scale models. Nucleic Acids Res. 44, D515–22 (2016).

9. J. R. Karr, J. C. Sanghvi, D. N. Macklin, M. V. Gutschow, J. M. Jacobs, B. Bolival Jr, N. Assad-Garcia, J. I. Glass, M. W. Covert, A whole-cell computational model predicts phenotype from genotype. Cell 150, 389–401 (2012).

10. Z. Ghaemi, J. R. Peterson, M. Gruebele, Z. Luthey-Schulten, An in-silico human cell model reveals the influence of spatial organization on RNA splicing. PLoS Comput. Biol. 16, e1007717 (2020).

11. D. N. Macklin, T. A. Ahn-Horst, H. Choi, N. A. Ruggero, J. Carrera, J. C. Mason, G. Sun, E. Agmon, M. M. DeFelice, I. Maayan, K. Lane, R. K. Spangler, T. E. Gillies, M. L. Paull, S. Akhter, S. R. Bray, D. S. Weaver, I. M. Keseler, P. D. Karp, et al., Simultaneous cross-evaluation of heterogeneous E. coli datasets via mechanistic simulation. Science 369, eaav3751 (2020).

12. R. S. Malik-Sheriff, M. Glont, T. V. N. Nguyen, BioModels—15 years of sharing computational models in life science. Nucleic Acids (2020).

13. E. Agmon, R. K. Spangler, A Multi-Scale Approach to Modeling E. coli Chemotaxis. Entropy 22 (2020).

14. J. Singla, K. M. McClary, K. L. White, F. Alber, A. Sali, R. C. Stevens, Opportunities and Challenges in Building a Spatiotemporal Multi-scale Model of the Human Pancreatic β Cell. Cell 173, 11–19 (2018).

15. Z. Fu, E. R. Gilbert, D. Liu, Regulation of insulin synthesis and secretion and pancreatic Beta-cell dysfunction in diabetes. Curr. Diabetes Rev. 9, 25–53 (2013).

16. B. de Finetti, Theory of Probability: A critical introductory treatment (John Wiley & Sons, 2017).

17. D. Koller, N. Friedman, F. Bach, Probabilistic Graphical Models: Principles and Techniques (MIT Press, 2009).

18. T. Baltru š aitis, C. Ahuja, L. Morency, Multimodal Machine Learning: A Survey and Taxonomy. IEEE Trans. Pattern Anal. Mach. Intell. 41, 423–443 (2019).

19. K. L. White, J. Singla, L. Valentina, J.-H. Chen, A. Ekman, L. Sun, X. Zhang, J. P. Francis, Li, W. Lin, K. Tseng, G. McDermott, F. Alber, A. Sali, C. Larabell, R. C. Stevens, Visualizing subcellular rearrangements in intact β-cells using soft X-ray tomography. Science Advances (2020).

20. C. Dalla Man, R. A. Rizza, C. Cobelli, Meal simulation model of the glucose-insulin system. IEEE Trans. Biomed. Eng. 54, 1740–1749 (2007).

21. D. J. Drucker, M. A. Nauck, The incretin system: glucagon-like peptide-1 receptor agonists and dipeptidyl peptidase-4 inhibitors in type 2 diabetes. Lancet 368, 1696–1705 (2006).

22. M. A. Nauck, J. J. Meier, Incretin hormones: Their role in health and disease. Diabetes Obes. Metab. 20 Suppl 1, 5–21 (2018).

23. S. S. Thazhath, C. S. Marathe, T. Wu, J. Chang, J. Khoo, P. Kuo, H. L. Checklin, M. J. Bound, R. S. Rigda, B. Crouch, K. L. Jones, M. Horowitz, C. K. Rayner, The glucagon-like peptide-1 (GLP-1) receptor agonist, exenatide, inhibits small intestinal motility, flow, transit and absorption of glucose in healthy subjects and patients with type 2 diabetes: a randomised controlled trial. Diabetes, db150893 (2015).

24. X. Zhu, R. Hu, M. Brissova, R. W. Stein, A. C. Powers, G. Gu, I. Kaverina, Microtubules Negatively Regulate Insulin Secretion in Pancreatic β Cells. Dev. Cell 34, 656–668 (2015).

25. F. M. Dekking, C. Kraaikamp, H. P. Lopuhaä, L. E. Meester, A Modern Introduction to Probability and Statistics: Understanding Why and How (Springer Science & Business Media, 2005).

26. H. F. Inman, E. L. Bradley, The overlapping coefficient as a measure of agreement between probability distributions and point estimation of the overlap of two normal densities. Communications in Statistics - Theory and Methods 18, 3851–3874 (1989).

27. M. A. Kurowski, J. M. Bujnicki, GeneSilico protein structure prediction meta-server. Nucleic Acids Res. 31, 3305–3307 (2003).

28. Y. Freund, R. E. Schapire, A Decision-Theoretic Generalization of On-Line Learning and an Application to Boosting. Journal of Computer and System Sciences 55, 119–139 (1997).

29. K. Lasker, M. Topf, A. Sali, H. J. Wolfson, Inferential optimization for simultaneous fitting of multiple components into a CryoEM map of their assembly. J. Mol. Biol. 388, 180–194 (2009).

30. A. Grover, A. Kapoor, E. Horvitz, A Deep Hybrid Model for Weather Forecasting in Proceedings of the 21th ACM SIGKDD International Conference on Knowledge Discovery and Data Mining, KDD ‘15., (Association for Computing Machinery, 2015), pp. 379–386.

31. T. L. Blundell, G. G. Dodson, D. Mercola, D. C. Hodgkin, The structure, chemistry and biological activity of insulin. AdvProteinChem 26, 279–402 (1972).

32. B. Szigeti, Y. D. Roth, J. A. P. Sekar, A. P. Goldberg, S. C. Pochiraju, J. R. Karr, A blueprint for human whole-cell modeling. Current Opinion in Systems Biology 7, 8–15 (2018).

33. D. S. Goodsell, M. A. Franzen, T. Herman, From Atoms to Cells: Using Mesoscale Landscapes to Construct Visual Narratives. J. Mol. Biol. 430, 3954–3968 (2018).

34. M. Feig, Y. Sugita, Whole-Cell Models and Simulations in Molecular Detail. Annu. Rev. Cell Dev. Biol. 35, 191–211 (2019).

35. J. Ma, M. K. Yu, S. Fong, K. Ono, E. Sage, B. Demchak, R. Sharan, T. Ideker, Using deep learning to model the hierarchical structure and function of a cell. Nat. Methods 15, 290 (2018).

36. A. J. Willsey, M. T. Morris, S. Wang, H. R. Willsey, N. Sun, N. Teerikorpi, T. B. Baum, G. Cagney, K. J. Bender, T. A. Desai, D. Srivastava, G. W. Davis, J. Doudna, E. Chang, V. Sohal, D. H. Lowenstein, H. Li, D. Agard, M. J. Keiser, et al., The Psychiatric Cell Map Initiative: A Convergent Systems Biological Approach to Illuminating Key Molecular Pathways in Neuropsychiatric Disorders. Cell 174, 505–520 (2018).

37. D. S. Tourigny, A. Goldberg, J. R. Karr, Simulating single-cell metabolism using a stochastic flux-balance analysis algorithm. bioRxiv (2020).

38. J. R. Stiles, D. Van Helden, T. M. Bartol, E. E. Salpeter, M. M. Salpeter, Miniature endplate current rise times< 100 s from improved dual recordings can be modeled with passive acetylcholine diffusion from a synaptic vesicle. Proceedings of the National Academy of Sciences 93, 5747–5752 (1996).

39. I. I. Moraru, J. C. Schaff, B. M. Slepchenko, M. L. Blinov, F. Morgan, A. Lakshminarayana, F. Gao, Y. Li, L. M. Loew, Virtual Cell modelling and simulation software environment. IET Syst. Biol. 2, 352–362 (2008).

40. A. E. Cowan, I. I. Moraru, J. C. Schaff, B. M. Slepchenko, L. M. Loew, Spatial modeling of cell signaling networks. Methods Cell Biol. 110, 195–221 (2012).

41. J.-J. Tapia, A. S. Saglam, J. Czech, R. Kuczewski, T. M. Bartol, T. J. Sejnowski, J. R. Faeder, MCell-R: A Particle-Resolution Network-Free Spatial Modeling Framework. Methods Mol. Biol. 1945, 203–229 (2019).

42. M. Tomita, K. Hashimoto, K. Takahashi, T. S. Shimizu, Y. Matsuzaki, F. Miyoshi, K. Saito, S. Tanida, K. Yugi, J. C. Venter, C. A. Hutchison 3rd, E-CELL: software environment for whole-cell simulation. Bioinformatics 15, 72–84 (1999).

43. G. T. Johnson, L. Autin, M. Al-Alusi, D. S. Goodsell, M. F. Sanner, A. J. Olson, cellPACK: a virtual mesoscope to model and visualize structural systems biology. Nat. Methods 12, 85–91 (2015).

44. I. Yu, T. Mori, T. Ando, R. Harada, J. Jung, Y. Sugita, M. Feig, Biomolecular interactions modulate macromolecular structure and dynamics in atomistic model of a bacterial cytoplasm. Elife 5 (2016).

45. R. Kalhor, H. Tjong, N. Jayathilaka, F. Alber, L. Chen, Genome architectures revealed by tethered chromosome conformation capture and population-based modeling. Nat. Biotechnol. 30, 90–98 (2011).

46. B. G. Wilhelm, S. Mandad, S. Truckenbrodt, K. Kröhnert, C. Schäfer, B. Rammner, S. J. Koo, G. A. Cla ß en, M. Krauss, V. Haucke, H. Urlaub, S. O. Rizzoli, Composition of isolated synaptic boutons reveals the amounts of vesicle trafficking proteins. Science 344, 1023–1028 (2014).

47. A. B. Noske, A. J. Costin, G. P. Morgan, B. J. Marsh, Expedited approaches to whole cell electron tomography and organelle mark-up in situ in high-pressure frozen pancreatic islets. J. Struct. Biol. 161, 298–313 (2008).

48. C. Ounkomol, S. Seshamani, M. M. Maleckar, F. Collman, G. R. Johnson, Label-free prediction of three-dimensional fluorescence images from transmitted-light microscopy. Nat. Methods 15, 917–920 (2018).

49. K. A. Gerbin, T. Grancharova, R. Donovan-Maiye, M. C. Hendershott, J. Brown, S. Q. Dinh, J. L. Gehring, M. Hirano, G. R. Johnson, A. Nath, A. Nelson, C. M. Roco, A. B. Rosenberg, M. Filip Sluzewski, M. P. Viana, C. Yan, R. J. Zaunbrecher, K. R. Cordes Metzler, V. Menon, et al., Cell states beyond transcriptomics: integrating structural organization and gene expression in hiPSC-derived cardiomyocytes. bioRxiv, 2020.05.26.081083 (2020).

50. D. P. Hoffman, G. Shtengel, C. S. Xu, K. R. Campbell, M. Freeman, L. Wang, D. E. Milkie, H. A. Pasolli, N. Iyer, J. A. Bogovic, D. R. Stabley, A. Shirinifard, S. Pang, D. Peale, K. Schaefer, W. Pomp, C.-L. Chang, J. Lippincott-Schwartz, T. Kirchhausen, et al., Correlative three-dimensional super-resolution and block-face electron microscopy of whole vitreously frozen cells. Science 367 (2020).

51. S. Calhoun, M. Korczynska, D. J. Wichelecki, B. San Francisco, S. Zhao, D. A. Rodionov, M. W. Vetting, N. F. Al-Obaidi, H. Lin, M. J. O’Meara, D. A. Scott, J. H. Morris, D. Russel, S. C. Almo, A. L. Osterman, J. A. Gerlt, M. P. Jacobson, B. K. Shoichet, A. Sali, Prediction of enzymatic pathways by integrative pathway mapping. Elife 7 (2018).

52. Q. Li, H. Tjong, X. Li, K. Gong, X. J. Zhou, I. Chiolo, F. Alber, The three-dimensional genome organization of Drosophila melanogaster through data integration. Genome Biol. 18, 145 (2017).

53. J.-K. Hériché, S. Alexander, J. Ellenberg, Integrating Imaging and Omics: Computational Methods and Challenges. Annu. Rev. Biomed. Data Sci. 2, 175–197 (2019).

54. A. Jha, M. R. Gazzara, Y. Barash, Integrative deep models for alternative splicing. Bioinformatics 33, i274–i282 (2017).

55. T. Stuart, A. Butler, P. Hoffman, C. Hafemeister, E. Papalexi, W. M. Mauck 3rd, Y. Hao, M. Stoeckius, P. Smibert, R. Satija, Comprehensive Integration of Single-Cell Data. Cell 177, 1888–1902.e21 (2019).

56. G. Lee, B. Kang, K. Nho, K.-A. Sohn, D. Kim, MildInt: Deep Learning-Based Multimodal Longitudinal Data Integration Framework. Front. Genet. 10, 617 (2019).

57. G. J. Félix-Mart ínez, J. R. God í nez-Fernández, Mathematical models of electrical activity of the pancreatic β-cell: a physiological review. Islets 6, e949195 (2014).

58. N. R. Johnston, R. K. Mitchell, E. Haythorne, M. P. Pessoa, F. Semplici, J. Ferrer, L. Piemonti, P. Marchetti, M. Bugliani, D. Bosco, E. Berishvili, P. Duncanson, M. Watkinson, J. Broichhagen, D. Trauner, G. A. Rutter, D. J. Hodson, Beta Cell Hubs Dictate Pancreatic Islet Responses to Glucose. Cell Metab. 24, 389–401 (2016).

59. D. Avrahami, A. Klochendler, Y. Dor, B. Glaser, Beta cell heterogeneity: an evolving concept. Diabetologia 60, 1363–1369 (2017).

60. M. S. Klemen, J. Dolen š ek, M. S. Rupnik, A. Sto ž er, The triggering pathway to insulin secretion: Functional similarities and differences between the human and the mouse β cells and their translational relevance. Islets 9, 109–139 (2017).

61. M. Orecná, R. Hafko, MZ. Bacová, J. Podskocová, D. Chorvát Jr, V. Strbák, Different secretory response of pancreatic islets and insulin secreting cell lines INS-1 and INS-1E to osmotic stimuli. Physiol. Res. 57, 935–945 (2008).

62. M. Skelin, M. Rupnik, A. Cencic, Pancreatic beta cell lines and their applications in diabetes mellitus research. ALTEX 27, 105–113 (2010).

63. P. Yang, Multi-Grid Method. Encyclopedia of Tribology, 2333–2339 (2013).

64. H. M. Berman, The Protein Data Bank: a historical perspective. Acta Crystallogr. A 64, 88–95 (2007).

65. S. K. Burley, G. Kurisu, J. L. Markley, H. Nakamura, S. Velankar, H. M. Berman, A. Sali, T. Schwede, J. Trewhella, PDB-Dev: a Prototype System for Depositing Integrative/Hybrid Structural Models. Structure 25, 1317–1318 (2017).

66. M. Hucka, A. Finney, B. J. Bornstein, S. M. Keating, B. E. Shapiro, J. Matthews, B. L. Kovitz, M. J. Schilstra, A. Funahashi, J. C. Doyle, H. Kitano, Evolving a lingua franca and associated software infrastructure for computational systems biology: the Systems Biology Markup Language (SBML) project. Syst. Biol. 1, 41–53 (2004).

67. D. Waltemath, J. R. Karr, F. T. Bergmann, V. Chelliah, M. Hucka, M. Krantz, W. Liebermeister, P. Mendes, C. J. Myers, P. Pir, B. Alaybeyoglu, N. K. Aranganathan, K. Baghalian, A. T. Bittig, P. E. P. Burke, M. Cantarelli, Y. H. Chew, R. S. Costa, J. Cursons, et al., Toward Community Standards and Software for Whole-Cell Modeling. IEEE Trans. Biomed. Eng. 63, 2007–2014 (2016).

68. J. R. Karr, J. C. Sanghvi, D. N. Macklin, A. Arora, M. W. Covert, WholeCellKB: model organism databases for comprehensive whole-cell models. Nucleic Acids Res. 41, D787–92 (2013).

69. R. K. Tripathy, I. Bilionis, Deep UQ: Learning deep neural network surrogate models for high dimensional uncertainty quantification. J. Comput. Phys. 375, 565–588 (2018).

70. A. Cozad, N. V. Sahinidis, D. C. Miller, Learning surrogate models for simulation-based optimization. AIChE J. 60, 2211–2227 (2014).

71. S. F. Sousa, P. A. Fernandes, M. J. Ramos, Protein-ligand docking: current status and future challenges. Proteins 65, 15–26 (2006).

72. H. M. Berman, P. D. Adams, A. A. Bonvin, S. K. Burley, B. Carragher, W. Chiu, F. DiMaio, T. E. Ferrin, M. J. Gabanyi, T. D. Goddard, P. R. Griffin, J. Haas, C. A. Hanke, J. C. Hoch, G. Hummer, G. Kurisu, C. L. Lawson, A. Leitner, J. L. Markley, et al., Federating Structural Models and Data: Outcomes from A Workshop on Archiving Integrative Structures. Structure 27, 1745–1759 (2019).

73. C. I. Byrnes, A. Isidori, New results and examples in nonlinear feedback stabilization. Syst. Control Lett. 12, 437–442 (1989).

74. M. S. German, Glucose sensing in pancreatic islet beta cells: the key role of glucokinase and the glycolytic intermediates. Proc. Natl. Acad. Sci. U. S. A. 90, 1781–1785 (1993).

75. P. E. MacDonald, J. W. Joseph, P. Rorsman, Glucose-sensing mechanisms in pancreatic β-cells. Philos. Trans. R. Soc. Lond. B Biol. Sci. 360, 2211–2225 (2005).

76. N. R. Gandasi, P. Yin, M. Omar-Hmeadi, E. Ottosson Laakso, P. Vikman, S. Barg, Glucose-Dependent Granule Docking Limits Insulin Secretion and Is Decreased in Human Type 2 Diabetes. Cell Metab. 27, 470–478.e4 (2018).

77. S. Costes, Targeting protein misfolding to protect pancreatic beta-cells in type 2 diabetes. Curr. Opin. Pharmacol. 43, 104–110 (2018).

78. O. Moltedo, P. Remondelli, G. Amodio, The Mitochondria–Endoplasmic Reticulum Contacts and Their Critical Role in Aging and Age-Associated Diseases. Frontiers in Cell and Developmental Biology 7 (2019).

79. M. Giacomello, L. Pellegrini, The coming of age of the mitochondria–ER contact: a matter of thickness. Cell Death Differ. 23, 1417–1427 (2016).

80. Zhongying Wang, Tatyana Gurlo, Aleksey V. Matveyenko, Peiyu Wang, Madeline Rosenberger, Jason A. Junge, Raymond C. Stevens, Kate L. White, Scott E. Fraser, Peter C. Butler, Live cell imaging of glucose-induced metabolic coupling of β and α cell metabolism in health and type 2 diabetes. in review (2020).

81. R. Bertram, L. S. Satin, A. S. Sherman, Closing in on the Mechanisms of Pulsatile Insulin Secretion. Diabetes 67, 351–359 (2018).

82. A. V. Matveyenko, D. Liuwantara, T. Gurlo, D. Kirakossian, C. Dalla Man, C. Cobelli, M. F. White, K. D. Copps, E. Volpi, S. Fujita, P. C. Butler, Pulsatile portal vein insulin delivery enhances hepatic insulin action and signaling. Diabetes 61, 2269–2279 (2012).

83. F. Sacco, S. J. Humphrey, J. Cox, M. Mischnik, A. Schulte, T. Klabunde, M. Schäfer, M. Mann, Glucose-regulated and drug-perturbed phosphoproteome reveals molecular mechanisms controlling insulin secretion. Nature Communications 7 (2016).

84. H. Tjong, W. Li, R. Kalhor, C. Dai, S. Hao, K. Gong, Y. Zhou, H. Li, X. J. Zhou, M. A. Le Gros, C. A. Larabell, L. Chen, F. Alber, Population-based 3D genome structure analysis reveals driving forces in spatial genome organization. Proc. Natl. Acad. Sci. U. S. A. 113, E1663–72 (2016).

85. C. Ionescu-Tirgoviste, P. A. Gagniuc, E. Gubceac, L. Mardare, I. Popescu, S. Dima, M. Militaru, A 3D map of the islet routes throughout the healthy human pancreas. Sci. Rep. 5, 14634 (2015).

86. A. Pisania, G. C. Weir, J. J. O’Neil, A. Omer, V. Tchipashvili, J. Lei, C. K. Colton, S. Bonner-Weir, Quantitative analysis of cell composition and purity of human pancreatic islet preparations. Lab. Invest. 90, 1661–1675 (2010).

87. L. Sun, A. Sali, Data-driven Brownian dynamics simulations: Application to insulin secretory pathway in pancreatic β-cells. to be submitted.

88. M. Kanehisa, S. Goto, KEGG: kyoto encyclopedia of genes and genomes. Nucleic Acids Res. 28, 27–30 (2000).

89. L. E. Fridlyand, N. Tamarina, L. H. Philipson, Bursting and calcium oscillations in pancreatic beta-cells: specific pacemakers for specific mechanisms. Am. J. Physiol. Endocrinol. Metab. 299, E517–32 (2010).

90. L. E. Fridlyand, N. Tamarina, L. H. Philipson, Modeling of Ca2+ flux in pancreatic beta-cells: role of the plasma membrane and intracellular stores. Am. J. Physiol. Endocrinol. Metab. 285, E138–54 (2003).

91. L. E. Fridlyand, L. Ma, L. H. Philipson, Adenine nucleotide regulation in pancreatic beta-cells: modeling of ATP/ADP-Ca2+ interactions. Am. J. Physiol. Endocrinol. Metab. 289, E839–48 (2005).

92. Q. Ni, A. Ganesan, N.-N. Aye-Han, X. Gao, M. D. Allen, A. Levchenko, J. Zhang, Signaling diversity of PKA achieved via a Ca2 -cAMP-PKA oscillatory circuit. Nature Chemical Biology 7, 34–40 (2011).

93. T. Redij, R. Chaudhari, Z. Li, X. Hua, Z. Li, Structural Modeling and in Silico Screening of Potential Small-Molecule Allosteric Agonists of a Glucagon-like Peptide 1 Receptor. ACS Omega 4, 961–970 (2019).

94. L. Marselli, M. Suleiman, M. Masini, D. Campani, M. Bugliani, F. Syed, L. Martino, D. Focosi, F. Scatena, F. Olimpico, F. Filipponi, P. Masiello, U. Boggi, P. Marchetti, Are we overestimating the loss of beta cells in type 2 diabetes? Diabetologia 57, 362–365 (2014).

95. M. Fehr, S. Lalonde, I. Lager, M. W. Wolff, W. B. Frommer, In vivo imaging of the dynamics of glucose uptake in the cytosol of COS-7 cells by fluorescent nanosensors. J. Biol. Chem. 278, 19127–19133 (2003).

